# Microscale physiological events on the human cortical surface

**DOI:** 10.1101/770743

**Authors:** Angelique C. Paulk, Jimmy C. Yang, Daniel R. Cleary, Daniel J. Soper, Milan Halgren, Alexandra R. O’Donnell, Sang Heon Lee, Mehran Ganji, Yun Goo Ro, Hongseok Oh, Lorraine Hossain, Jihwan Lee, Youngbin Tchoe, Nicholas Rogers, Kivilcim Kiliç, Sang Baek Ryu, Seung Woo Lee, John Hermiz, Vikash Gilja, István Ulbert, Daniel Fabó, Orrin Devinsky, Joseph R. Madsen, Donald L. Schomer, Emad N. Eskandar, Jong Woo Lee, Douglas Maus, Anna Devor, Shelley I. Fried, Pamela S. Jones, Brian V. Nahed, Sharona Ben-Haim, Sarah K. Bick, Robert Mark Richardson, Ahmed M. T. Raslan, Dominic A. Siler, Daniel P. Cahill, Ziv M. Williams, G. Rees Cosgrove, Shadi A. Dayeh, Sydney S. Cash

## Abstract

Despite ongoing advancements in our understanding of the local single-cellular and network-level activity of neuronal populations in the human brain, extraordinarily little is known about their ‘intermediate’ microscale local circuit dynamics. Here, we utilized ultrahigh density microelectrode arrays and a rare opportunity to perform intracranial recordings across multiple cortical areas in human participants to discover three distinct classes of cortical activity that are not locked to ongoing natural brain rhythmic activity. The first included fast waveforms similar to extracellular single unit activity. The other two types were discrete events with slower waveform dynamics and were found preferentially in upper layers of the grey matter. They were also observed in rodents, non-human primates, and semi-chronic recordings in humans via laminar and Utah array microelectrodes. The rates of all three events were selectively modulated by auditory and electrical stimuli, pharmacological manipulation, and cold saline application and had small causal co-occurrences. These results suggest that with the proper combination of high resolution microelectrodes and analytic techniques it is possible to capture neuronal dynamics that lay between somatic action potentials and aggregate population activity and that understanding these intermediate microscale dynamics may reveal important details of the full circuit behavior in human cognition.

## Introduction

Using neural recordings to uncover how the brain and underlying circuitry support behavior has traditionally concentrated on two ends of the spectrum visible with extracellular neural recordings: single-cell spiking activity or local field potentials (LFPs) arising from the summation of synaptic activity in large populations of neurons. However, other forms of neural electrical signals such as sodium or calcium potentials in the dendritic arbor, back-propagating action potentials, axonal action potentials, ephaptic signaling, and more can be captured using intracellular recordings from single cells or optical imaging (e.g. 2-photon calcium imaging). These other forms of neural electrical signals have, of course, been the focus of neuroscience investigation but, due to technical limitations, have been difficult to study in vivo with large-scale circuit analysis, particularly in humans. Significant advances in imaging and recording in rodents have begun to come close (Abdelfattah et al. 2019; Yildirim et al. 2019) and advances in microelectrode array technology have made exploring neural activity at the microscale level increasingly accessible (Jia et al. 2019). Yet, investigations of these other forms of activity outside single cell extracellular spikes or LFP oscillations, however, have not been performed in vivo in humans.

We took advantage of the development of high density poly(3,4-ethylenedioxythiophene) polystyrene sulfonate (PEDOT:PSS) microelectrodes to detect microscale neural activity from large areas of the cortical surface in humans, rodents, and non-human primates (Wilks et al. 2011; Sessolo et al. 2013; Khodagholy et al. 2015, 2016, 2017; Cellot et al. 2016; Hermiz et al. 2020). PEDOT:PSS microelectrodes represent a next stage in technologies developed for recording microscale neuronal activity in humans which also include laminar arrays of microelectrodes (Ulbert, Halgren, et al. 2001; Ulbert, Karmos, et al. 2001; Ulbert et al. 2004; Wang et al. 2005; Fabó et al. 2008; Cash et al. 2009; Keller et al. 2010; Halgren et al. 2018, 2019) and 2D arrays of penetrating microelectrodes (the Utah array, (Nordhausen et al. 1994; Hochberg et al. 2006; Schevon et al. 2008, 2010; Truccolo et al. 2008; Waziri et al. 2009; Keller et al. 2010). Importantly, specialized recording (including novel microelectrodes) and analytic approaches have been used to explore axonal action potentials (Robbins et al. 2013; Stuart and Spruston 2015; Jun et al. 2017), dendritic calcium spikes (Ross 2015; Suzuki and Larkum 2017; Golding et al. 2018), and backpropagating action potentials (Moore et al. 2017; Jia et al. 2019) in rodents. However, a systematic study of these events across multiple species including humans under multiple conditions has yet to be undertaken. We took advantage of high density capabilities of PEDOT:PSS arrays and other microelectrode systems to detect and examine microscale cortical activity beyond action potentials and LFP oscillations in humans, rodents, and a non-human primate.

We performed recordings using PEDOT:PSS microelectrodes from patients undergoing craniotomies for the surgical resection of tumor or epilepsy focus. We also analyzed recordings from semi-chronic placement of laminar microelectrodes and Utah arrays. We observed both LFP dynamics (typical of pial recordings with metal contacts; (Kaiju et al. 2017)) and high frequency events similar to the single unit activity seen in intraparenchymal recordings with penetrating metal electrodes (Khodagholy et al. 2015, 2016). What was remarkable, though, is that we noted two other classes of morphologically, spatially, and temporally distinct events with features between single unit activity and slower oscillatory events which we call Type 2 and Type 3 events. We then performed a series of tests to validate these events are not just artifact and examine these events across multiple microelectrode types and multiple species with several analyses to finally conclude that these events have a physiological origin in brain activity and are not a part of ongoing spontaneous oscillations.

## Materials and Methods

### Human Participant Recordings in the Operating Room

Recordings from humans in the operating room were acquired with 37 participants (mean=39.7 years old, ranging from 22 to 62; 18 females; all but 7 individuals right handed **Supplementary Tables 1-2**) who were scheduled for surgical resection of cortical tissue as a result of tumor or epilepsy at Massachusetts General Hospital (MGH), Brigham and Women’s Hospital (BWH), University of California San Diego (UCSD), and Oregon Health and Science University (OHSU). Patients were already scheduled for a craniotomy for concurrent clinical intraoperative neurophysiological monitoring or testing for mapping motor, language, and sensory regions and removal of tissue.

In some cases (N=9), pro-convulsant medications (methohexital or alfentanil) were given intravenously to the participants in the process of clinically mapping the putative seizure focus while recording with the clinical electrodes. This choice was made by the clinical team. In another subset of cases (N=8), cold saline was applied for the purposes of reducing overactive tissue clinically or to determine how underlying activity changes with a change in temperature. Finally, with some participants (N=11), repetitive sounds (1 sec between each sound) were played back to the participants via a speaker in the operating room (OR; N=4) or via headphones to the participant (N=7) using Presentation software (Neurobehavioral Systems) to determine if there was an auditory response to the sounds. The different types of sounds were either real words, nonsense syllables which were not English words, and noise-vocoded sounds (which involved a white noise convolution of the real words (Travis et al. 2013; Hermiz et al. 2018; Kaestner 2018).

Following the surgery, the preoperative T1-weighted MRI was reconstructed using FreeSurfer scripts (Reuter et al. 2010, 2012; Dykstra et al. 2012) (http://surfer.nmr.mgh.harvard.edu). The reconstructions were then aligned to images obtained during surgery using blender software (https://www.blender.org/) and MMVT (Holmes et al. 1998; Dykstra et al. 2012; Felsenstein and Peled 2017; Peled et al. 2017). The method involved projecting the surgical image onto the patient’s reconstructed brain using blender and then placing a 3D model of the PEDOT:PSS electrode array on that location similar to other coregistration approaches (Holmes et al. 1998; Postelnicu et al. 2009; Dykstra et al. 2012) (**Fig. 1; Supplementary Fig. 1-3**; **Supplementary Fig. 1-3**).

**Figure 1.**
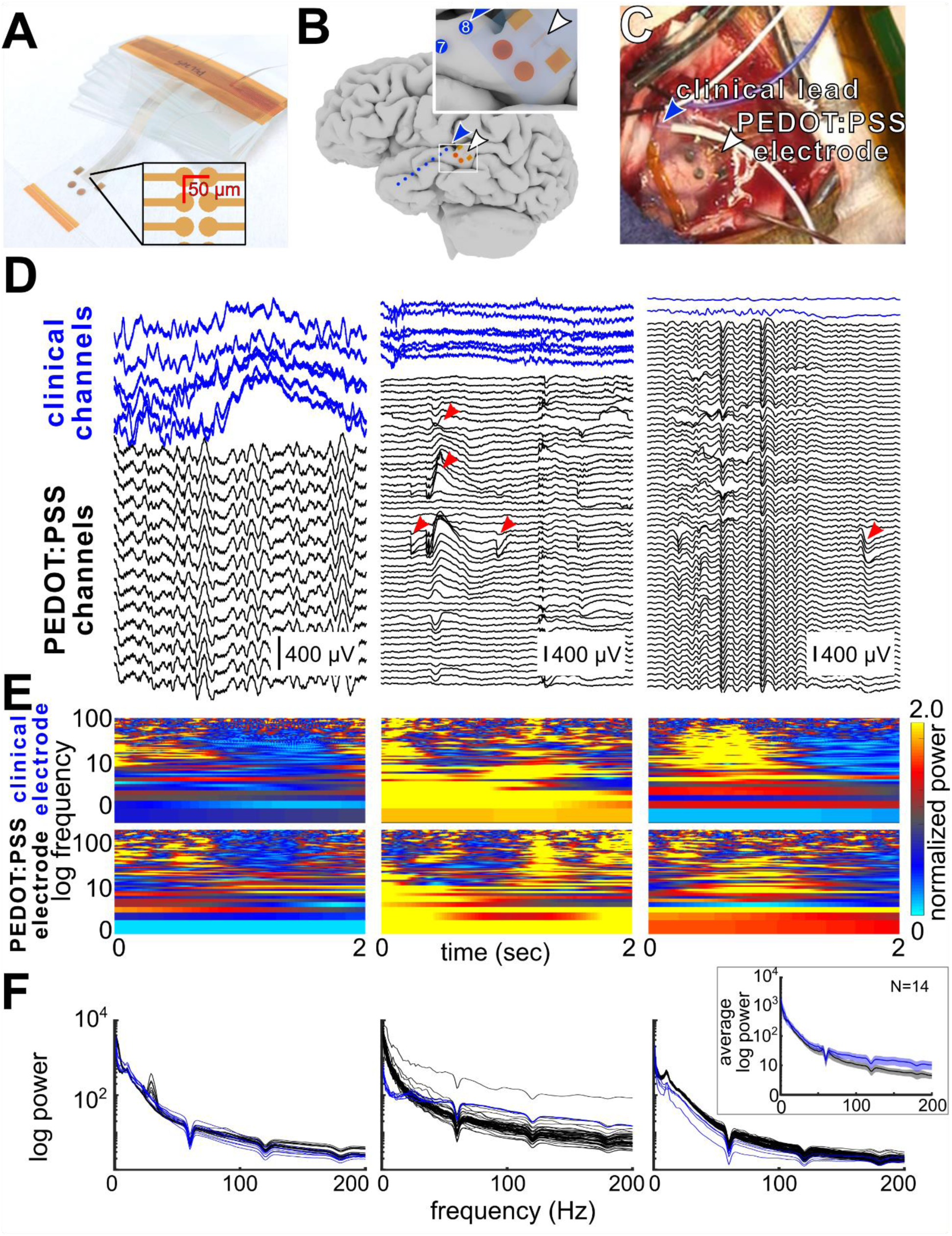
PEDOT:PSS recording set up, fast events, and spectrum. **A**. PEDOT:PSS 2-column 128 channel grid electrode layout (two rows of 64 channels separated by 50 µm) and dimensions. **B**. Example of relative positioning. The larger circles and squares on the electrode array are large PEDOT contacts to allow for orientation of the small electrode contacts (the 2-column grid) in between the two large squares. The inset square is a zoomed in view of the PEDOT:PSS electrode. **C**. Photograph of microelectrode (white arrowhead) and clinical array (blue arrowhead) on the human cortical surface alongside reference electrodes (white and purple wire). **D**. Example broadband recordings from three participants (IP03, IP15, and IP18) of clinical leads (blue lines) and the PEDOT:PSS grid (black lines, bottom), with unitary events (red arrowheads, grid middle, and right traces. The geometry of the PEDOT:PSS grid in all three recordings was a 128 channel 2-column grid extending 3150 µm × 50 µm with a 50 µm pitch between sites. Time scale for traces in **D** is the same as in **E**. **E**. Normalized power spectrograms of simultaneous individual clinical and PEDOT:PSS and one same channel trace shown in **D**. Each column of LFP traces in **D** match the spectrograms shown in **E**. **F**. Individual average power spectra curves per PEDOT:PSS channel (black) and clinical channels (blue) for IP03, IP15, and IP18. Inset: Average values across participants (N=14). Shaded regions indicate s.e.m.

### Semi-chronic Human Participant Recordings in the Epilepsy Monitoring Unit

Implantations of the laminar and Utah array electrodes were performed on patients with pharmacologically resistant epilepsy undergoing surgery to locate and resect seizure foci (**Supplementary Table 3**). All decisions concerning placement were made with clinical input, particularly since both the Utah arrays and laminar microelectrodes were inserted into cortex likely to be resected. Laminar recordings were made in the United States and Hungary while the Utah array recordings were performed in the United States. The electrodes were implanted for a period of days to 2 weeks.

Nine patients (mean=23.375 years old, ranging from 12 to 42; 4 females) were implanted with laminar electrodes. Laminar microelectrodes were implanted perpendicular to the cortex in noneloquent tissue that was ultimately resected and has been reported on previously (Ulbert, Halgren, et al. 2001) and also used in a number of studies (Ulbert, Karmos, et al. 2001; Ulbert et al. 2004; Wang et al. 2005; Fabó et al. 2008; Cash et al. 2009; Keller et al. 2010; Halgren et al. 2018, 2019). Each laminar probe spanned the cortical depth with a length of 3.5 mm and diameter of 0.35 mm. Each array was comprised of 24 electrodes in a single row with diameters of 40mm spaced at spaced between 175 μm center to center. Neural data was recorded and stored continuously with 2 kS/s/channel rate in the EEG range simultaneously with MUA range activity sampled at 20 kS/s/channel. Please see those prior publications for methodological details.

A further eight patients (mean=32 years old, ranging from 21 to 47; 2 females) were implanted with Utah array (also called a NeuroPort array) electrodes (Blackrock Microsystems, USA), which has been used in several previous studies (Hochberg et al. 2006; Truccolo et al. 2008; Waziri et al. 2009; Schevon et al. 2010). The 4mm × 4mm microelectrode array is composed of 100 platinum-tipped silicon probes at two different depths. In four cases, the electrodes penetrated 1.0mm into the cortex. In another four cases, the electrodes penetrated 1.5 mm into cortex.

### Ethics statement

All patients voluntarily participated after fully informed consent according to NIH guidelines as monitored by the Partners Institutional Review Board (IRB) which provides coverage for Massachusetts General Hospital (MGH), Brigham and Women’s Hospital (BWH) and Boston Children’s Hospital (CH), by the UC San Diego Health IRB which provides coverage for University of California San Diego (UCSD), the New York University Medical Center IRB which provides coverage for New York University Medical Center, Committee on Clinical Investigations of the Beth Israel Deaconess Medical Center which provides coverage for Beth Israel Deaconess Medical Center (BIDMC), the Hungarian Medical Scientific Council, and the Oregon Health Sciences University IRB which provides coverage for the Oregon Health Sciences University (OHSU). Participants, whether recording in the operating room or with semi-chronic electrodes in the epilepsy monitoring unit, were informed that participation in the experiment would not alter their clinical treatment in any way, and that they could withdraw at any time without jeopardizing their clinical care.

### Non-Human Primate Surgical Details and Recording Methods

Intraoperative, intracranial neurophysiology recordings were acquired from one adult male rhesus macaque (*Macaca mulatta*, age 14; **Supplementary Table 4**) according to the Guide to the Care and Use of Laboratory Animals. All efforts were made to minimize discomfort, and the Institutional Animal Care and Use Committee at the Massachusetts General Hospital monitored care and approved all procedures. The macaque was placed under general endotracheal anesthesia (isoflurane) and placed into a stereotactic frame (Kopf; Kujunga, CA). A craniotomy over the visual cortex were performed using standard anatomic landmarks, and cortex was carefully exposed. Using gyral anatomy and vasculature over V1 versus V4, the V4 area was identified visually and a PEDOT:PSS electrode was placed over the region. Signals were recorded using the ORH128 Intan Recording System (Hermiz et al. 2016, 2018). The data was acquired at 30 kHz and filtered by default Intan setting with cutoffs 1 Hz to 7.5 kHz and using OpenEphys acquisition graphic-user interface software (Siegle et al. 2017) (http://www.open-ephys.org/), with the impedance tests of the electrodes during the experiments carried out using the Intan RHD2000 software from Intan Technologies (Los Angeles, CA). Data was extracted and processed using MATLAB (Mathworks, Natick, MA).

### Area V1 Mouse Surgical Details and Recording Methods

The care and use of mice (2-6 months old; C57BL/6J; Jackson Laboratory, Bar Harbor, ME; **Supplementary Table 4**) followed all federal and institutional guidelines, and the Institutional Animal Care and Use Committees of the Massachusetts General Hospital. Mice were deeply anaesthetized with a cocktail of injected ketamine hydrochloride (100 mg/kg intraperitoneal injection) and xylazine (10 mg/kg i.p. injection; N=1) or isoflurane administered as a gas prior and during the start of surgery (N=2). In the mouse given ketamine, additional ketamine (1/10 initial dose) was supplemented every 30 min to maintain the plane of anesthesia.

Anesthetized mice were placed into a stereotaxic frame (Narishige, Japan) for the craniotomy as well as all subsequent testing. A heating blanket on the floor of the frame was used to maintain body temperature at 37°C. A craniotomy was performed on 4.5 mm × 4.5 mm area around the primary visual cortex (V1) defined by a stereotaxic coordinate (AP : −3.8, ML: −2 mm; (Paxinos and Franklin 2001)). A small craniotomy was made in the skull, the PEDOT:PSS array was placed over the exposed visual cortex over intact dura. Once the PEDOT:PSS array was positioned on V1, neural signals were recorded using the ORH128 Intan Recording System, acquiring data at 30 KHz and filtered by the default Intan setting (cutoffs 1 Hz to 7.5 kHz). Data was extracted and processed using MATLAB.

### Barrel Cortex (S1) Mouse Surgical Details and Recording Methods

ICR mice weighing 25-35 g were used in the experiments. All procedures were in accordance with a protocol approved by the Institutional Animal Care and Use Committees of UC San Diego (protocol S07360; **Supplementary Table 4**). The mice were anesthetized with isoflurane, a femoral artery was catheterized to allow recording of blood pressure, and a tracheotomy was performed for ventilation of the mice. A metal headpost was fixed to the skull using dental acrylic, a craniotomy and durotomy were performed over the right whisker barrel and surrounding cortex. The exposure was surrounded by a dental acrylic well in order to keep it filled with artificial CSF until the electrode array was placed. The exposure was dried prior to the electrode placement, and then once the array was implanted it was covered with 0.7% agarose in artificial CSF. The electrodes arrays used were arranged in square grids with 0.2 mm spacing, and 50 µm diameter contacts. The reference electrode was silver-chloride ball placed between muscle tissue exposed for the craniotomy. Prior to recording the mice were administered pancuronium and artificially ventilated, and prior to stimulus trials the mice were switched from isoflurane to alpha-chloralose anesthesia. Whisker flick stimuli were presented every 2 seconds though analyses and detections of events were performed on baseline, non-stimulated periods of time. Neural signals were recorded at 20 kHz using the Intan recording system.

### PEDOT:PSS electrode Device Fabrication

The fabrication of the PEDOT:PSS device is similar to previously established protocols with slight modifications as noted below (Uguz et al. 2016; Ganji, Atsunori, et al. 2017; Ganji, Kaestner, et al. 2017). Si wafers in batches of 6 wafers were used as substrate carriers for deposition of parylene C layers. Following substrate cleaning, the diluted Micro-90 (0.1%) – an anti-adhesion layer – was spun-cast at 1300 rpm on the substrate and an NR9-6000PY negative resist was used for metal electrode definition on the parylene C layers. O_2_ plasma (Plasma Etch PE100) was applied for 2 min (150W RF power, 5 sccm O_2_) for descum step to remove residual photoresist in patterned regions prior to the evaporation of a 15 nm Cr adhesion layer and a 100 nm Au contact layer on top of the 4 μm thick parylene C layer (epidural mouse recording) or 10 μm thick parylene C layer (subdural recordings for mouse, NHP, and humans). After lift-off, an O_2_ plasma (Plasma Etch PE100) was applied for 2 min (150W RF power, 5 sccm O_2_) to activate the surface of parylene C for enhancing the adhesion of the subsequent 2 μm thick encapsulating parylene C layer. Another higher concentrated Micro-90 (1% as opposed to 0.1% for the first layer) layer was spun cast and a third parylene C layer (≈ 1.9–2.5 μm) was deposited as sacrificial layer. To define the patterns on electrode sites, a 10 μm thick 2010 SU-8 photoresist layer was exposed and developed using Karl Suss MA6 mask aligner and SU-8 developer. Prior to PEDOT:PSS deposition, O_2_ plasma (Oxford Plasmalab 80, 200 W RF power, 32 min) was used to etch the openings in the third and second parylene C layers all the way to the contact sites. Additionally, a hole at the center of the circular array was etched through all parylene C layers. The remainder of the fabrication process remained essentially the same as previously reported (Rivnay et al. 2016; Uguz et al. 2016; Ganji, Kaestner, et al. 2017).

Four different PEDOT:PSS electrode designs were used. One electrode type involved 2-columns of 64-channel 20 µm diameter contact sites with 50 µm center to center pitch between adjacent contacts, which we call the 50 µm pitch 2-column grid (N=25; **Fig. 1; Supplementary Fig. 1-2; 4**). A second array had the same material make-up but the center-to-center pitch was 800 µm, which we call the 800 µm pitch 2-column grid (N=1; **Supplementary Fig. 3**, **5**). A third array was a 128-channel grid with concentric rings at varying distances between each electrode site, which we call the circular grid ∼4 mm in diameter (N=11; **Supplementary Fig. 3**, **5**). Finally, one array was used while recording over the mouse barrel cortex which was a square grid of 32 channels.

**Figure 2.**
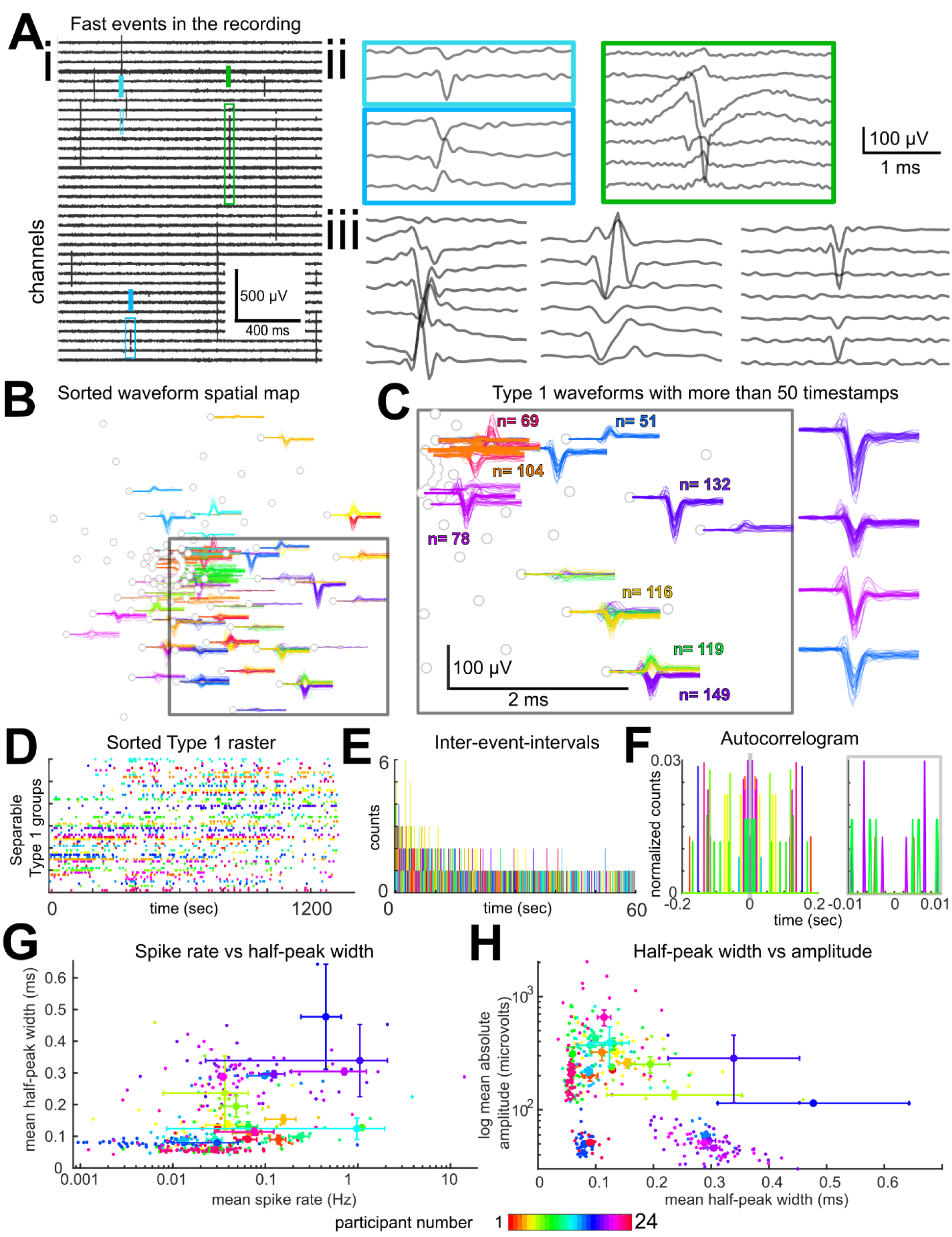
Fast, Type 1 events recorded from the surface of the human cortex. **A**. **i**-**iii**. Example recording from participant IP37. In **ii**, the events in colored boxes from **i** are shown at higher resolution. In **iii**, further examples from the rest of the recording are shown with spatial spread across multiple channels and voltage field reversals. Scale bar in **ii** applies to the voltage data shown in **ii** and **iii**. **B**, **C.** Example fast waveforms from the same participant, mapped to the circular grid locations across the PEDOT:PSS microelectrode array. The waveforms are color coded corresponding to different clusters as determined in Kilosort using spatial locations of waveforms on the PEDOT:PSS grids combined with spike times in a model (see **Materials and Methods**). The grey dots indicate good channels with low impedances on the circular PEDOT:PSS array. **D-F**. Single example recordings from the same clusters as in **B** and **C** shown as a raster plot across the microelectrode grid, as inter-event intervals (IEIs, **E**) and auto-correlograms (**F**) for a subset of the clusters with different time scales at ±0.2 sec (left plot) and ±0.01 sec (right plot). **G, H**. Comparisons of half-peak widths vs event frequencies (spike rate; **H**) and half-peak widths versus trough to peak amplitudes (**G**) of the sorted waveforms were compared across all participants. N=24. Error bars indicate s.e.m.

**Figure 3.**
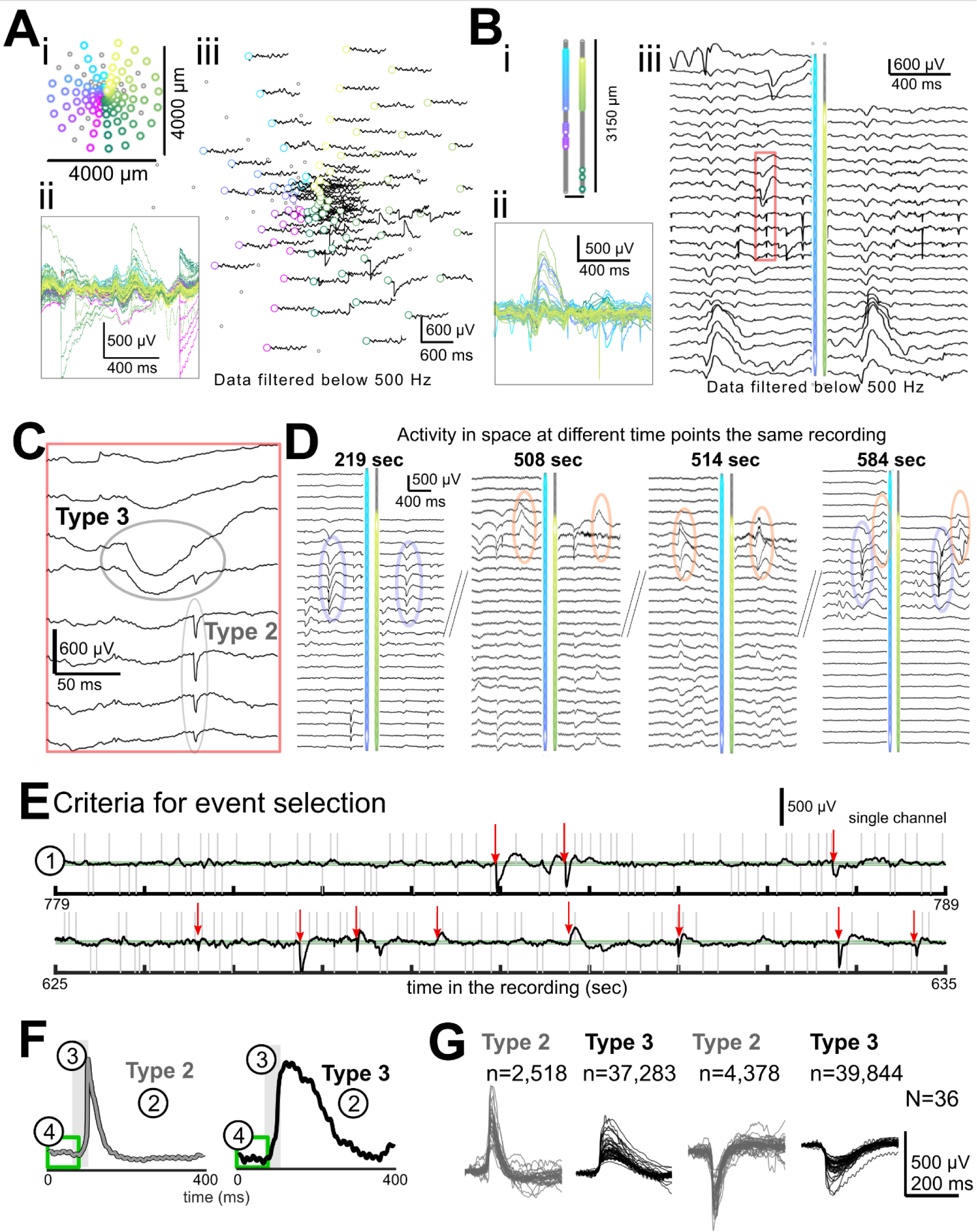
Classifiable events can be identified across recordings and participants. **A**. Recording from participant IP25 in time (**Ai-ii**) and mapped to the circular grid (**Aiii**). Waveform color-coding matches channel color-coding with cyan to magenta (channels 1-64) and green to yellow (channels 65-128) indicating the four separate 32 channel amplifier banks used in the recording (Bank A: 1-32; Bank B: 33-64; Bank C: 65-96; Bank D: 97-128). Only the channels which passed criteria are shown as non-grey dots. **B**. Example recording (electrodes which passed criteria represented as non-grey dots, **Bi**) from participant IP15 showing activity changes in time (**Bii**) and a proportion of electrodes and their activity mapped to the two-column electrode array (**Biii**). **C**. Zoomed view of the red box in **Biii** showing Type 2 and 3 events. **D**. Example recordings of events at different time points (for IP15) showing only a subset of electrode channels. Repeated events through time for two different event examples are shown (orange and blue ovals). **E**-**F**. Stepwise criteria to detect Type 2 and 3 events, with template waveforms shown in **F**. (1) The first step is to detect absolute peaks that are greater than 25 µV, as shown in example traces from a single channel and case (IP15). Grey vertical lines are detected cross-threshold peaks. Green lines are the thresholds. Red arrows are the events which fulfill all four for selection criteria. (2) The next step is detecting peaks that correlate with the templates ≥ 0.80. (3) Selected events had a second derivative at onset which is greater than 2 in the time period at onset, indicated by the shaded grey box. Finally, (4) waves were kept if they had no large deflections before the event onset, i.e., within 100 ms prior to crossing the 25 µV threshold, indicated by the green boxes. **G**. Average waveforms per participant (N=36) for the different classes of events. All voltage data shown were low pass filtered using the high cut-off frequency of 500 Hz.

### Data Acquisition: Acute Intraoperative Recordings

Most human intracranial recordings in the OR from the PEDOT:PSS electrodes were made using a custom Intan recording system with a sampling rate of either 20 kHz or 30 kHz (filtered from 1 Hz to 7500 Hz to reduce aliasing; Intan Corporation, Los Angeles, CA). During acquisition, recordings were referenced to sterile ground and recording reference needle electrodes (Medtronic) placed in nearby muscle tissue (often scalp) as deemed safe by the neurosurgeon. After the electrode array was laid on tissue and the grounds placed, we tested impedances using the Intan system using the RHD2000 series software (Intan Corp.). In most cases, we used OpenEphys (Siegle et al. 2017), an open-source software which allowed us to save and visualize activity across all 128 channels as well as the analog input. Trigger signals produced by a custom MATLAB program indicating the hour, minute, and second were additionally sent to both the analog input for the Intan system as well as to the TRIG input to the clinical Quantum system (Natus) which was acquiring the concurrent clinical electrode recordings. The clinical Quantum system recorded neural activity at either 4096 Hz (at MGH) or 512 Hz (at BWH). At UCSD and OHSU, we recorded neural activity using the PEDOT:PSS electrode only. To test whether activity we observed was due to the Intan system, we also recorded neural PEDOT:PSS activity using a second recording system with a sampling rate of 30 kHz using a high impedance headstage and amplifiers input to a Neural Signal Processor (Blackrock Microsystems, USA). In one case, two PEDOT:PSS electrodes (one connected to the Intan system and the other to the Blackrock system) were used in the same case for simultaneous recording for a side-by-side comparison (**Supplementary Fig. 3**).

### Data Acquisition: Semi-Chronic Laminar and Utah Array Recordings

For the laminar recordings, differential recordings were made from each pair of successive contacts. After wideband filtering (DC-10 kHz), the signal was split into a low frequency field potential band (filtered at 0.2–500 Hz, gain 1000x, digitized at 2 kHz, 16 bit) and stored continuously. Periods of time where there were clear artifacts were rejected from the laminar recordings.

Utah array recordings were made from 96 active electrodes and data were sampled at 30 kHz per electrode (0.3–7 kHz bandwidth) using Blackrock hardware. Recording data from the Utah arrays was taken from 30 minutes of baseline activity while the participants were awake (around 11 am) 3-4 days after implant.

### Saline Recordings

While in the OR in a subset of cases (N=6), we performed several minutes of saline bath recordings before and after recording from the brain using the same equipment and electrodes as connected in the recordings on neural tissue, minutes either before or after recording from human brain. For all comparisons, we performed the same analyses and classification steps as listed below on these saline tests to determine if we could observe the same events as identifying in the brain OR recordings. In two cases, we purposefully moved the electrode in the saline to reproduce some of the same movements observed in the case.

### Electrical stimulation

Electrical stimulation was performed in N=11 cases (**Supplementary Table 1-2**). In most cases, neural stimulation was carried out for clinical purposes such as performing motor or language mapping (Mueller and Morris 1993; Borchers et al. 2012). Stimulation was either performed using a Prass Standard Monopolar Stimulating Probe (motor mapping cases, Medtronic, USA), Disposable Double Ball Tip Nerve Stimulating Probe (speech mapping cases, Caldwell, USA), or the Nicolet Cortical Stimulator for Neurosurgery (Natus Neurology Inc., Middleton, WI, USA). For the clinical stimulation mapping (N=9), trains (at 60 Hz) or pulses of stimulation of variable length (ranging from .1 to 1 sec) and amplitude (ranging from 1 to 10 mA) were delivered via a hand-held bipolar wand at different locations based on the surgical flow of the case. Research stimulation (N=2) consisted of train stimulation (100 Hz, 0.5 sec duration) delivered at 5 second intervals via a pair of surface clinical strip electrodes near the PEDOT:PSS electrode.

### Intraoperative Video monitoring

With the patient’s informed consent, we recorded video of the surgery. The video recordings were acquired by a camera set in the operating room light (Black Diamond Video). We approximated the timing of the video recordings by annotating in the test when the electrode was placed on and removed from the brain which could also be seen in the recordings. We additionally annotated the timing and location of stimulation via bipolar contacts or movements of the PEDOT:PSS electrode in the field to confirm the timing of noisy signals relative to additional experimental events. Pictures of the preparation were also taken to illustrate the electrode location (**Fig. 1**).

### Data Analysis

Data analysis was performed using custom programs in MATLAB. Cross-correlation between resampled analog trigger signals was used to align the triggers between multiple systems, both between the neural data and between the audio recordings. Since the recordings were brief (4-20 minutes) and electrodes could be moved at any point, we used the video and audio recordings as well as the voltage recordings to determine periods of stable recordings.

‘Raw’ voltage signals were taken directly from the Intan recordings. LFP data were decimated to 1000Hz and demeaned relative to the entire recording. Multi-unit activity (MUA) or spike-related activity was detected by high pass filtering the signal above 250 Hz. Channels with excessive line noise, had high impedances (>100 kΩ), or which fluctuated across the voltage range were removed from the analysis. We also noted that, since these recordings are necessarily transient in the operating room, we would find some channels would become noisier during the recordings over time as others. For this reason, we performed root mean square (RMS) calculations of electrodes for the ‘quiet’ periods of time when we identified when the electrode was not being moved and when there were no visible inter-ictal discharges (IIDs) or stimulation (**Supplementary Fig. 6**). We found that, by and large, the recordings in channels that had within-range impedances also had low RMS values (around or below an RMS of 5). However, there were a subset of channels that had high RMS values (closer to an RMS of 20-50) which we removed from further analyses. These channels tended to have recordings which were unstable through time based on the fluctuations in voltage and the presence of 60 Hz noise. After excluding the recordings from one participant, 48.5 ± 23.26 % of the electrode contacts (channels) survived channel rejection criteria across patients (**Supplementary Table 2; Supplementary Fig. 6;** see **Materials and Methods**). We found, as we progressed in cases and we gained experience, the % of good electrode contacts improved to closer to 80%. We also identified the locations of channels which survived the rejection criteria and found that the locations good channels varied considerably across the grids (**Supplementary Fig. 4-5**) and did not correspond to amplifier bank or physical connectivity (e.g. particular cables or connectors). **Type 1 Sorting**

We first sorted Type 1 waveforms from the data high pass filtered above 250 Hz using a Butterworth filter via Offline Sorter (v4, Plexon, Inc., Texas, USA). We identified unit waveforms as activity within the range of 15-500 µV with a duration between 500-1500 µsec. However, with the pitch and high density of the electrodes, we found that we were identifying too many ‘units’ since the putative units likely spanned multiple channels in the recording. For this reason, we used Kilosort (Pachitariu et al. 2016). Kilosort identifies clusters of waveforms as repeated events based on the spatial arrangement of the electrode as well as waveform morphology which allowed us to identify putative repeated events with similar waveform shapes across the recording as is done with typical single unit spike sorting using high spatial resolution electrodes such as Neuropixel (Jun et al. 2017; Jia et al. 2019). Kilosort omits spike detection and PCA and instead combines template waveforms, associated spike locations and spike times into one model. Before processing, we only applied Kilosort to data which had no movement or stimulation artifact and removed noisy channels. Putative event clusters (the equivalent to single units) were then rejected if the captured waveforms as sampled looked to be artifact (such as with stimulation) or if the units had highly regular or too short inter-event-intervals. Rejecting events with inter-event intervals (IEIs) corresponding with 60 Hz noise or which occurred at the same time as the stimulation, we also identified the fast waveforms in the raw and original high pass filtered data to confirm their timing and channel location. Considering the low event count, we also attempted to merge clusters based on the spatial mapping of the recordings. However, when we did merge clusters, we found significantly more violations of the minimum refractory period of 2-5 ms (meaning the clusters were more likely multi-unit activity when merged) indicating to us that these events needed to be separated to represent activity more similar to single unit activity. Therefore, we did not merge clusters nor did we use autocorrelations of event times to make any decisions to merge clusters. Of note is that we could not easily use signal to noise ratio (SNR) as a method to test the quality of these clusters as SNR usually assumes detecting events on one or a few channels. In this case, we are identifying clusters of events using Kilosort with a high spatial resolution across 7-10 channels on the PEDOT:PSS grid where a spatial and waveform characteristics define each cluster across the spatial map. This approach can lead to large waveforms on some channels and no waveforms on other channels (**Fig. 2**). Therefore, SNR, and SNR variance, may not be as informative compared to other recording approaches.

### Type 2 and 3 Event Detection and Template Matching

Since other ‘events’ involved both short (Type 1) and long (traveling LFP wave) timescales and varied in size and directions of deflection (due to changes in voltage polarity across electrodes), we performed a first-pass visualization to detect and annotate electrode movement, stimulation, or the presence of inter-ictal discharges (IIDs) using Wonambi, a visualization tool originally used in sleep scoring (https://wonambi-python.github.io/). We rejected any time periods when the electrodes were being moved in the recording. We identified IIDs for these analyses as events of ∼250 ms duration and similar to previously published IID waveforms (Curtis et al. 2012; Janca et al. 2015) but did not include those IID waveforms in the subsequent event classification analyses since we considered those events outside of the scope of the current study.

Next, we detected peaks larger than 25 µV and took 400 ms snippets of data around each event (−100 ms to 300 ms) per channel. Among those events, we then re-aligned events relative to the largest rising or falling edge (for easier comparisons across waveforms). After examining all the waveforms both singly and using PCA (**Supplementary Fig. 8**), we found we could further remove noise by performing template matching to the latter two of three main waveform types: 1) Type 1: fast repeated waveform events, detected between 250 and 6000 Hz; 2) Type 2: peaks with a sharp rise and slow fall, ∼10-30 ms absolute half-peak duration, detected below 500 Hz; or 3) Type 3: peaks with a slow rise and slow fall, ∼100 ms absolute half-peak duration, detected below 500 Hz. Templates were constructed from average identified waveforms across the data set based on the rates of the events and most common waveforms identified through PCA, which included the Type 2 and 3 events (**Supplementary Fig. 9**). Since we were performing the template matching on the LFP, not the raw data, we would not be able to detect Type 1 using this approach. Correlations to each waveform above 0.8 were used to exclusively bin each event into one of these three types (**Supplementary Fig. 9**).

Another possible explanation for these events is that they result from electrical noise or some kind of mechanical artifact. To address this, we examined saline recordings performed in the operating room using the same electrodes for several minutes before and after recording from the brain in N=6 cases (**Supplementary Fig. 10-12**). In two cases, we applied movement to the saline bowl in an attempt to reproduce the waveforms observed in the neural recordings. First, the Type 1 events observed in the neural recordings were not present in any of the saline recordings (data not shown). We either never or very rarely detected Type 2 events in any of the saline recordings, with or without movement in saline (**Supplementary Fig. 10-12**). When we moved the electrode in the saline, waveforms similar to the Type 3 events could be identified in the saline recordings, although, these were rare compared with those of the neural recordings, and the waveforms were visibly slower to rise and fall, resulting in a lower second derivative of the onset time (**Supplementary Fig. 10**), without the sharp initial rise of Type 3 waveforms observed in the neural recordings which meant that brain-recorded Type 3 waveforms had higher second derivatives around the waveform onset (**Supplementary Fig. 10-12**). After examining the saline recording detections and finding overlap between on brain vs in saline recordings in the waveforms following our detection approach, we found we needed to add another detection step which involved thresholding based on the second derivative at ± 50 ms around the onset of the waveform (**Supplementary Fig. 10-12**). Finally, as a step to remove the possibility of including ongoing oscillatory waveforms, we added a final fourth step where we kept only waveforms with average absolute voltages < 25 µV in the preceding 100 ms before the event onset.

Even after these added waveform onset thresholding steps, most Type 2 and 3 events recorded on the surface of the brain survived the thresholding per participant (**Supplementary Fig. 10**). We found that both events happened significantly more frequently on brain tissue than in saline (p=0.0038; Kruskal-Wallis multiple comparison test; **Supplementary Fig. 12**), with a sum total of 59 Type 2 events and 854 Type 3 events across nine saline recordings and over 29 minutes versus 3,553 Type 2 events and 33,608 Type 3 events over brain tissue using the same electrodes over the same amount of time. Additionally, we found that 90% of events in saline spanned a single channel, with only 6 Type 2 events and 77 Type 3 events spanning multiple channels using our spatial extent detection window over a total period of 29 minutes (**Supplementary Fig. 12**).

The Type 2 and 3 classifications of LFP events were then mapped to the recordings in time to determine if they vary in rates, size, and temporal dynamics with the introduction of auditory stimuli, pro-convulsant medication, cold saline, or electrical stimulation. Within these categories, we quantified their rates in time as the inter-event interval (IEI) and the absolute amplitudes of the voltage deflections from baseline. We examined the rise and fall times relative to the peaks by quantifying the RC time constants τ_rise time_ and τ_fall time_ as in the following equation: ΔV = (V_final_ – V_start_)*(1-1/e^(time/ τ)), where τ_rise time_ was calculated as time it took for the voltage to reach 0.63 of the maximum peak and τ_fall time_ was calculated as the time it took for the voltage to reach 0.37 of the fall back to baseline.

Additional measures included the temporal and spatial spread of the events in a recording which included identifying events which were concurrent in time within 5 ms of each other across channels and calculating their spatial spread based on the electrode layout. Peri-stimulus time histograms (PSTHs) involved binning activity in time around specific events, such as the application of pro-convulsant medications, cold saline, auditory stimulation, or electrical stimulation. The bins were one at 100 ms or at other time ranges depending on the treatment. PSTHs and inter-event interval counts were normalized relative to baseline by dividing each bin by the average bins during the baseline recordings before averaging PSTHs across participants. Cross-covariance measures were calculated by converting the timing of the Type 1, Type 2, and Type 3 events in into 1 ms bins throughout the recording (at a sample rate of 1000 Hz) and then performing a cross-covariance between the event types. Autocorrelograms were performed by a leave-out method which involved taking the timestamp per detected event and cross-correlating that timestamp in that recording with all other events in that Type 1 group or per channel for the Type 2 and 3 events. This should result in a lack of a peak at time = 0 with patterns of rhythmicity emerging if there are periodic elements.

### Spectral Analyses

Power spectral analyses to compare the simultaneously recorded clinical and PEDOT:PSS recordings were performed using 5 minutes of ‘quiet’ periods of data (without movement, stimulation, or tasks and before application of medication or cold saline) by taking the real value of the Morlet wavelet coefficient (power) at a 1-Hz spectral resolution using a moving window of 0.5 sec moving every 10 ms, calculated using the Fieldtrip toolbox (www.ru.nl/fcdonders/fieldtrip). This approach allowed us to observe dynamics through time and as an average.

Event-triggered spectral analyses were performed by taking the 30 seconds around each event (whether the Type 1, 2, or 3 events) of the data sampled at 1000 Hz and then calculating power again by taking the real value of the Morlet wavelet coefficient (power) at a 1-Hz spectral resolution using a moving window of 0.5 sec moving every 10 ms, calculated using the Fieldtrip toolbox (www.ru.nl/fcdonders/fieldtrip) (Oostenveld et al. 2011). We then normalized the power at each frequency band by dividing each time step in power per frequency band by the mean power across all windows of time per frequency band. Finally, we averaged the power per the following frequency bands: 0-4, 4-8, 8-15, 15-30, 30-55, and 65-200 Hz.

As the Type 2 and 3 events had sharp onsets and had broadband effects on power, we clipped out the 200 ms around each actual event. We then performed two different spectral calculations to determine if the Type 2 and 3 events were part of an ongoing oscillation. One analysis involved a series of steps, which involved a) calculating spectrograms 10 sec before and after the Type 2/3 events (not including+/− 200 ms around each event); b) performing the same calculation but, instead of centering it around the Type 2/3 events, add or subtract a time in the range of +/− 10 seconds (jittered time-locked power calculation); c) calculating the difference between the two spectrograms on a per-channel basis and averaging per patient; and d) determining if that difference is significantly different from zero across the data set.

To measure the phase relationships between the same-channel LFP and the onset of the Type 2 and 3 events, we filtered the signal to each frequency band with eegfilt (EEGLab package; (Delorme and Makeig 2004)) and then calculated phase per frequency band (Hilbert transform) and averaged those phase values using the CircStats toolbox (Berens 2009). As we were performing these calculations on a per-channel and then per-case basis, we had to average phase values when the number of events per recording was greater than 5 as phase is highly susceptible to low event counts.

### Statistical analysis

All statistical comparisons were performed using non-parametric measures, so we did not test for normality. We tested comparisons with the Kruskal–Wallis test for non-equivalence of multiple medians followed by the use of the *post hoc* Tukey-Kramer method to determine statistically separable groups. For all comparisons, we first calculated the mean values per type of neural activity on the per-participant level and then comparing the medians between conditions. Measurements of activity were separated into baseline, during pharmacological manipulation, stimulation (auditory or electrical), or physiological manipulation. This resulted in measurements being taken from distinct samples for the manipulations aside from baseline activity, a period of time which was measured and compared multiply when needed to identify changes in activity. We used the Wilcoxon rank-sum test (two-sided) for comparisons between individual medians. We tested if values were significantly different from zero using the Wilcoxon signed rank test. We corrected by adjusting the target p-value (0.05) with a Bonferroni correction for the number of comparisons being done. For phase relationships, we performed the Rayleigh test for unipolar distributions (Berens 2009).

### Data and code availability

Open source acquisition software, OpenEphys (http://www.open-ephys.org/) and open source waveform sorting software Kilosort (https://github.com/cortex-lab/KiloSort) was used to record and analyze the neural data. Custom Matlab code (version R2019a) in combination with open source code from the Fieldtrip toolbox (http://www.fieldtriptoolbox.org/) was used for the majority of the analyses and is available upon request.

Custom Matlab code for detecting Type 2 and 3 events are available on Github (https://github.com/Center-For-Neurotechnology/LFP-Type2-and-Type3-Detection). The code requires the input of 1000 Hz neural data in microvolts and can be used to detect Type 2 and 3 events across the recording. Example data is provided along with plotting examples showing the resulting detections. Further data that support the findings of this study are available from the corresponding author upon reasonable request.

## Results

### PEDOT:PSS electrodes record neural activity from the surface of the human brain

We recorded neural activity from the cortical surface using PEDOT:PSS microelectrodes and clinical surface electrodes in patients undergoing craniotomy for either the removal of tumors (N=15) or for seizure focus resection for intractable epilepsy (N=22). PEDOT:PSS electrodes were 30 µm in diameter and had a center-to-center pitch between 50 and 800 µm. We used four different electrode designs (including circular, 2-column, and square grid designs) to cover areas ranging from 0.1575 to 40 mm^2^ (see **Materials and Methods**; **Fig. 1A-C**; **Supplemental Fig 1-5**). These different designs afforded different coverage over the cortex (**Supplemental Fig 1-5**). Fabricated on parylene C (5-14 um thickness) into either 32 or 128 channel arrays (**Supplementary Fig. 2-3**), electrodes were placed directly on and conformed to the pial surface of the exposed cortex. Recordings ranged from 4-25 minutes. With one participant (Intraoperative Participant 14, or IP14), the recordings were unstable due to electrode movement throughout the session and were therefore removed from subsequent analyses. Once the electrodes were placed on the brain, we tested their impedance values. For electrodes and contacts that passed our criteria (which include within range impedances and recording stability; see **Materials and Methods, Supplementary Fig. 4-6**), average impedance values were 44.5 ± 26.0 (mean ± SD) kOhm.

Next, we compared neural activity from the PEDOT:PSS recordings with concurrently recorded clinical electrodes (3 mm diameter platinum contact grids; **Fig. 1C**; N=14). The frequency spectrum of the ongoing electrical activity and recognizable events such as epileptiform inter-ictal discharges were similar between the microelectrodes and the clinical contacts (**Fig. 1D-E**). To determine if the oscillations captured by the PEDOT:PSS electrodes were similar to those recorded by the clinical contacts, we calculated power spectral density and spectrograms for both recording modalities from 1 to 200 Hz (**Fig. 1F**). Averaging the power spectra through spontaneous activity recordings, the spectra were not significantly different between the clinical contacts and the PEDOT:PSS recordings (p>0.07 for comparisons at each frequency step; Wilcoxon rank-sum test; N=14). We observed concurrent oscillations in both the PEDOT:PSS recordings and the clinical recordings. However, notably, the PEDOT:PSS recordings showed localized stand-alone events which were not detected by the clinical contacts (red arrowheads in **Fig. 1D**; **Supplementary Video 1**).

### PEDOT:PSS electrodes record putative spiking activity from the surface of the human brain

We detected a series of different ‘fast’ waveforms in recordings with PEDOT:PSS electrodes filtered between 250 and 6000 Hz. Previous studies with these devices have demonstrated recordings of single action potentials from the pial surface of various preparations, including in humans (Wilks et al. 2011; Sessolo et al. 2013; Khodagholy et al. 2015, 2016, 2017; Cellot et al. 2016; Kaiju et al. 2017; Hermiz et al. 2020). Some of these events (which we call Type 1 events) were similar in kinetics to single units (i.e., individual spikes) recorded with extracellular electrodes including prior surface PEDOT:PSS electrode recordings ((Khodagholy et al. 2015, 2016, 2017) **Fig. 2A**). We used the Kilosort software package (Pachitariu et al. 2016) to sort waveforms into a number of separable repeated events using a template matching approach that considers waveform spatial spread across electrode sites, arriving at clusters of repeated Type 1 events in 24 out of 36 participants (**Fig. 2**). We hypothesize that the presence or absence of Type 1 events was related to the contact with the cortical surface (which can vary from case to case per clinical considerations and variability in placement) as well as overlap over or near vessels or, in some cases, tumor tissue (see **Supplementary Material**). The average frequency (number of a given event / time in a given recording) across participants was 0.13 ± 0.75 with a median of 0.026 Hz (**Fig. 2C, 2D**). Inter-event interval (IEI) histograms as well as auto-correlograms showed diverse patterns within a recording (**Fig. 2E**, **F**) and across recordings (**Supplementary Fig. 7**). We did not observe consistent rhythmicity for these Type 1 events.

The sorted waveforms were generally negatively deflecting, though each event could be seen as negative on some channels and positive on the others (**Fig. 2**), with voltage amplitudes of 123.3 ± 89.57 µV and a median of 93.2 µV (averaged per cluster and then averaged across participants, resulting in 18,513 detected events subdivided into 420 identified, sortable, replicable ‘units’ detected across 24 participants, with 17.5 ± 16.8 clusters detected per participant; **Fig. 2G**, **2H**). The half peak widths for the largest waveforms per cluster across the data set averaged 0.49 ± 0.217 msec (**Fig. 2B**, **2C**, **2G**, **2H**). While fast compared to most somatic action potentials, these values are within the range for waveforms established by previous recordings using sharp electrodes (e.g. Utah array) in the human cortex or in rodents and non-human primates (Bartho et al. 2004; Keller et al. 2010; Vigneswaran et al. 2011; Robbins et al. 2013; Chan et al. 2014; Anastassiou et al. 2015; Hamilton et al. 2018; Trainito et al. 2019). However, some events had waveform kinetics faster or slower than what may be typical and amplitudes that were sometimes larger than what might be expected at the pial surface for neurons (**Fig. 2B**, **2C**, **2G**, **2H**).

### Slower microscale events are also detectable from the surface of the human brain

PEDOT:PSS electrode recordings also revealed microscale phenomena distinct from the fast waveform activity represented by Type-1 events (**Fig. 1, 2; Supplemental Video 1**). These discrete, slower events occurred across multiple channels in a single recording, varied in polarity across channels and in time, and could repeat through time across neighboring channels. Notably, these events could be separated into distinct classes based on duration, with some lasting 100-200 ms and others less than 20 ms. Such events were not restricted to a single amplifier chip or bank, cable, recording system, or other physical structure increasing the likelihood of their physiological origin (see **Materials and Methods**). Having observed these waveforms repeatedly across participants and recordings, we set out to classify and characterize the corresponding waveforms, seeking to determine whether these events were discrete spatially and temporally, could be a part of ongoing oscillations, could be modulated, or had a significant relationship to the detected fast waveforms or to each other.

Therefore, in addition to Type 1 (fast 1-2 ms duration events, detected in recordings filtered between 250 and 6000 Hz; **Fig. 1**), we defined two other types of events: Type 2 events with a sharp rise and slow fall, ∼10-30 ms absolute half-peak duration, detected in recordings filtered below 500 Hz and Type 3 events with a slow rise and slow fall, ∼100 ms absolute half-peak duration, detected in recordings filtered below 500 Hz (**Fig. 3A-F; Supplemental Video 1**). Initially, this classification was based on visual inspection of the waveforms and spectral analysis. However, we used PCA combined with k-means clustering to identify the most common baseline waveform templates for putative Type 2 and 3 events (**Supplementary Fig. 8**). These waveform templates became the basis of a template matching algorithm to detect and identify individual Type 2 and Type 3 events throughout the recordings. The criteria for identified Type 2 and 3 events were: 1) absolute waveforms that were >25 µV; 2) detected waveforms had a correlation above 0.8 with the template event waveform; 3) the second derivative at the onset of the waveform was greater than 2; and 4) the absolute average voltage in the 100 ms preceding the event onset was less than 25 µV (to reduce the chances of capturing ongoing oscillations; **Fig. 3E**, **F**; see **Materials and Methods**). The code for using these criteria to detect such events in other recordings is available for use by the community (https://github.com/Center-For-Neurotechnology/LFP-Type2-and-Type3-Detection).

If only to confirm that that our template-matching approach can both detect and separate Type 2 and 3 events, we found the differences in average time constant rise time (τ_rise time_) and time constant decay time (τ_fall time_) were significantly different between event types (following averaging values per patient and then comparing between types; N=36; p<0.00001; Wilcoxon rank-sum test; **Supplementary Fig. 9**). Type 2 events had an average time τ_rise time_ of 14.18 ± 8.42 ms and average τ_fall time_ of 30.62 ± 6.33 ms as averaged across participants (after taking the absolute voltage values, see **Materials and Methods**). In contrast, Type 3 events had τ_rise time_ of 28.62 ± 8.80 ms and τ_fall time_ of 81.48 ± 11.91 ms. More importantly, Type 3 events were significantly more frequent than Type 2 events (p<0.0001; Wilcoxon rank-sum test; **Fig. 3G; Supplementary Fig. 9**), with Type 3 events (n=55,406) occurring ∼9 times more often than Type 2 events (n=6,404) across recordings and participants. To demonstrate that these events repeated in time and space, we calculated the variance of the detected Type 2 and 3 events within and across channels to determine waveform consistency through time on the same channels. The idea is that, if there is less variance in these events on the per-channel level than across channels, the argument is that these events are repeated on the same channel. The voltage variance across channels was larger than within channels (**Supplementary Fig. 9**). While this difference was not significant (p>0.05, Wilcoxon rank-sum test, **Supplementary Fig. 9**), the use of template matching combined with the criteria to detect events limited how variable the detected waveforms could be between and across channels.

In multiple participants, we confirmed the presence of Type 2 and Type 3 events in various brain regions by sampling across brain regions, between cases and, in 12 instances, when moving the electrode to different brain areas during the recording (**Supplementary Fig. 1-3, 5**). There were no significant differences among recordings in the temporal lobe, dorsolateral prefrontal cortex, motor area (precentral gyrus), and the parietal lobe (p>0.3320; Kruskal-Wallis multiple comparison test). We did not observe significant differences in waveforms between the epilepsy and tumor surgery contexts or between awake and anesthetized cases (p>0.05; Wilcoxon rank-sum test; **Supplementary Fig. 1-2** and **Supplementary Fig. 9**). We found we could detect these events using different recording systems with the same electrodes (see **Materials and Methods**). In addition, we found almost no Type 1-3 events when recording over putative tumor tissue or when we flipped the array upside down such that the electrode pads were not facing the pial surface (N=1 in each case, see **Materials and Methods**) or in saline (**Supplementary Fig. 10-12**).

### Event Types 1, 2 and 3 have different spatial characteristics

Type 2 and 3 events spanned multiple channels (**Fig. 1, 2, 3**). We examined the spatial spread of concurrently Type 2 and 3 events with onset times that occurred within a 10 ms time window (chosen based on inter-event interval distributions; **Fig. 4A-B**). In windows with multiple channels detecting one event, we measured the average number of reporting channels, distance in space from the first to the last reporting channel (in µm), the area covered (in µm^2^), the speed of waveform travel (in m/sec), and the time span of the event (in sec). All these parameters were not significantly different between Type 2 and Type 3, though multichannel Type 3 events tended to cover more area and longer spatial distances (p>0.07; Wilcoxon rank-sum test; **Fig. 4B**). Using this approach, we found, that Type 2 and Type 3 events were observed, on average, across 5-10 channels across the entire array. This was equivalent to or slightly more than a 100 µm × 100 µm area, suggesting that these events are relatively localized (**Fig. 4B**).

**Figure 4.**
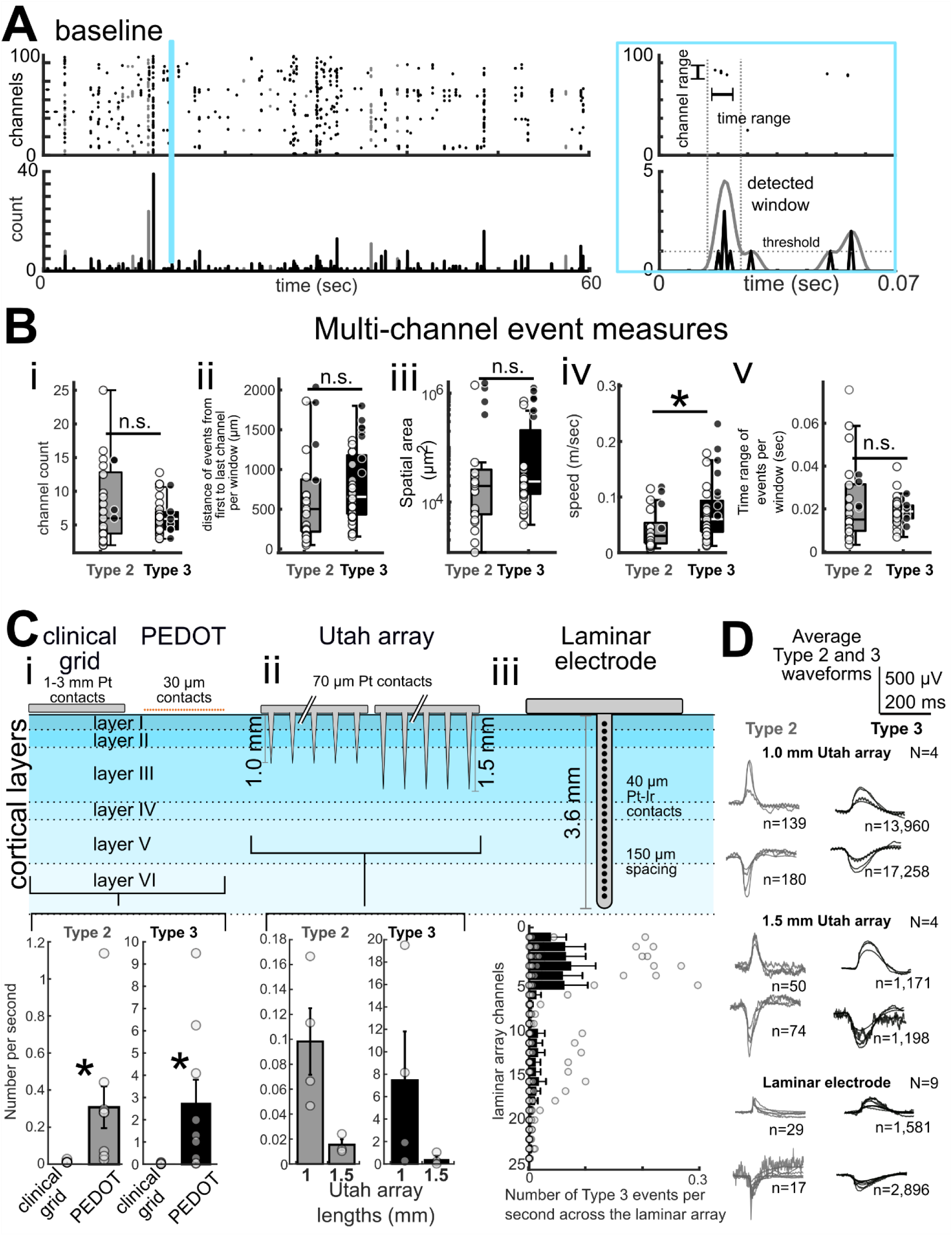
Temporal and spatial spread of Type 2 and 3 events. **A**. Example recording from participant IP17 illustrating methods for determining event spatial spread with the area in blue on the left expanded to show a zoomed in version of the time scale on the right. **Bi-v**. In cases where multiple channels detected one event, we compared the detected waveforms across channels, the time span covered by the event (sec), area covered (µm^2^), speed of the event propagation across channels (m/sec), and the distance travelled across the electrode array (µm). Box plots and confidence intervals around the mean values per measure per participant are indicated by whisker bars; center line indicates median, N=36. White dots are the 2-column grids, while black dots are the circular grid recordings per participant. Asterisks indicate p<0.05, Wilcoxon rank-sum test. **Ci-iii**. Top: Diagrams of the clinical surface electrodes, PEDOT:PSS array, Utah laminar arrays (1.0 and 1.5 mm depths), and laminar arrays showing relative sampling geometry of the different systems. Bottom: **Ci**. Significantly more Type 2 and 3 events per second occur in the PEDOT:PSS array recordings than in the clinical recordings (p<0.001; N=9; Wilcoxon rank-sum test). **Cii**. Significantly more Type 2 and 3 events are detected per second in the shallower Utah array recordings than the deeper recordings (p<0.001; N=4 each Utah array depth; Wilcoxon rank-sum test). **Ciii**. More Type 3 events are detected per second in superficial laminar contacts as evidenced by the average negative correlation between the channel number and Type 3 counts across the array (rho= −0.48 ± 0.38; significantly different to zero; p = 0.0313; Wilcoxon signed rank test; N= 9). **D.** Example average detected Type 2 and 3 events per participant with n=number of detected events.

**Figure 5.**
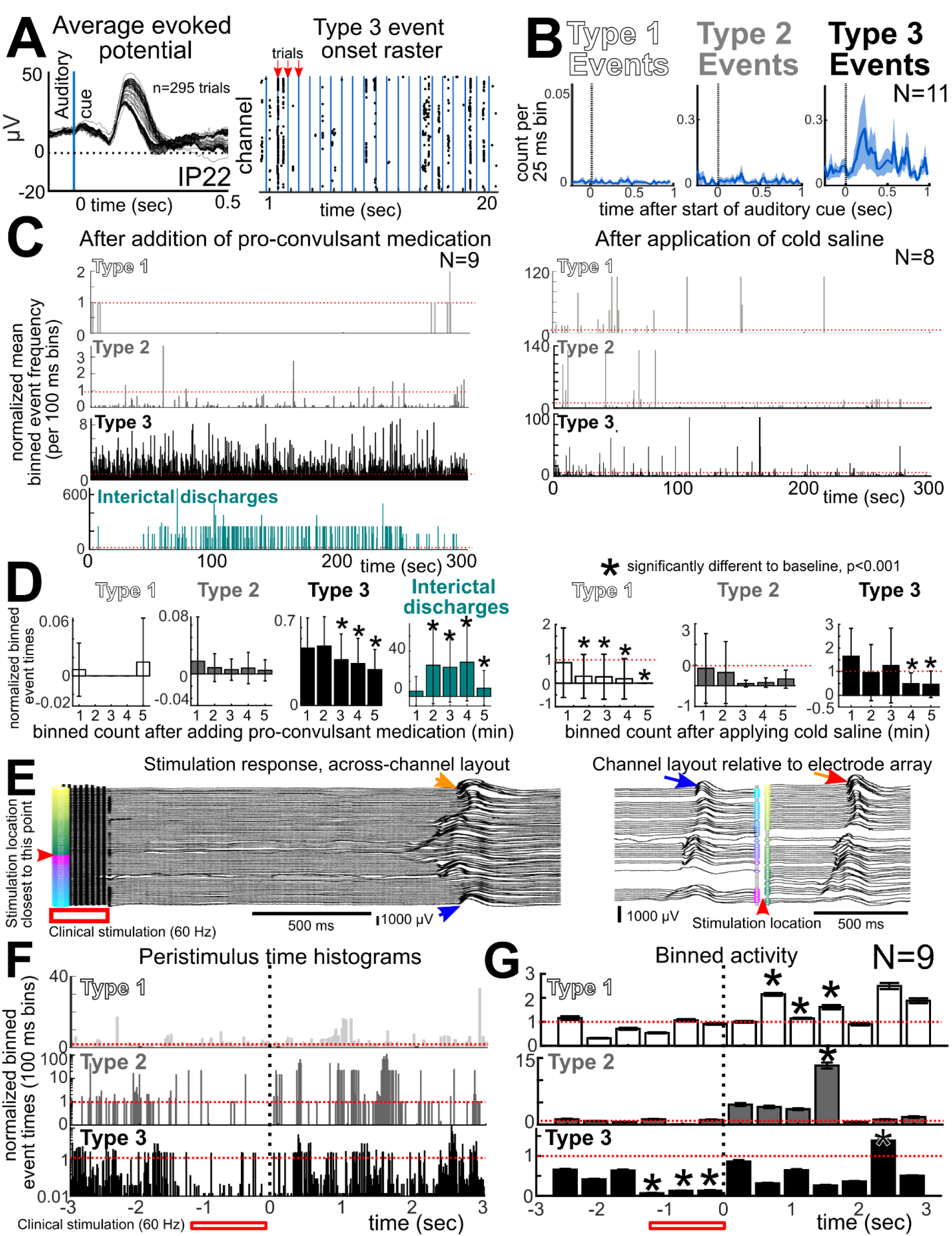
Manipulations alter rates of Type 1-3 events. **A.** Effects of auditory stimuli on Type 2-3 events. Left: Recording from participant IP22 showing neural dynamics during repeated auditory stimuli in the averaged evoked potentials across trials. Right: Raster plots of Type 3 events occurring across all trials with auditory stimulus trial times indicated by blue vertical lines. **B**. Left: Peri-stimulus time histograms (PSTHs) of each event type binned per 25 ms and then averaged across all patients (N=11). For the line plots, shaded error regions are all s.e.m. **C**. Effects of pro-convulsant medication and cold saline on Types 1 – 3 events and inter-ictal discharges. PSTH of binned activity following the administration of pro-convulsant medication (methohexital or alfentanil; left) or the application of cold saline (right) averaged across patients. Type 1 (top), Type 2 (middle), Type 3 events (middle) and IIDs (bottom) were binned every 100 ms and normalized relative to baseline by dividing each bin by the average baseline activity. The dotted red line is the baseline normalization value. **D**. Histogram of activity changes with 5-minute bins. Error bars are s.e.m., N=9 for pro-convulsant medication, N=8 for cold saline application. Asterisks indicate significant difference from baseline with p<0.001, Wilcoxon rank-sum test. **E**. Electrical stimulation changes in the rate of Type 1-3 events. Recording from participant IP07 showing a wave of activity (Type 3 event) following a a train of stimuli delivered at 60 Hz for 1-2 seconds stimulation during clinical mapping, shown in with the channels laid out along a single line (left) and time-voltage curves mapped relative to the 2-column grid electrode layout (activity in space, right). Blue and orange arrowheads indicate the same pair of channels as shown to the left time-voltage plots versus the right time-voltage plots relative to the electrode layout. The red arrowhead indicates the direction of stimulation relative to the electrode layout, though the bipolar stimulation was 1-2 cm away. **F**. Peri-stimulus time histograms of binned activity around the time of stimulation (bar below figure) averaged across patients (N=9, n>5 trials per participant). Type 1 (top), Type 2 (middle), and Type 3 (bottom) events were binned every 100 ms and normalized relative to the 1 sec before stimulation. The dotted red line is the normalized baseline value. Note: the y-axis is in log scaled increments. **G**. Histograms of activity in 0.5 sec bins before and after stimulation. p-values reflect differences between each 0.5 sec bin per event type, N=9; asterisks indicate significant difference from baseline with p<0.001, Wilcoxon rank-sum test. Error bars are all s.e.m.

**Figure 6.**
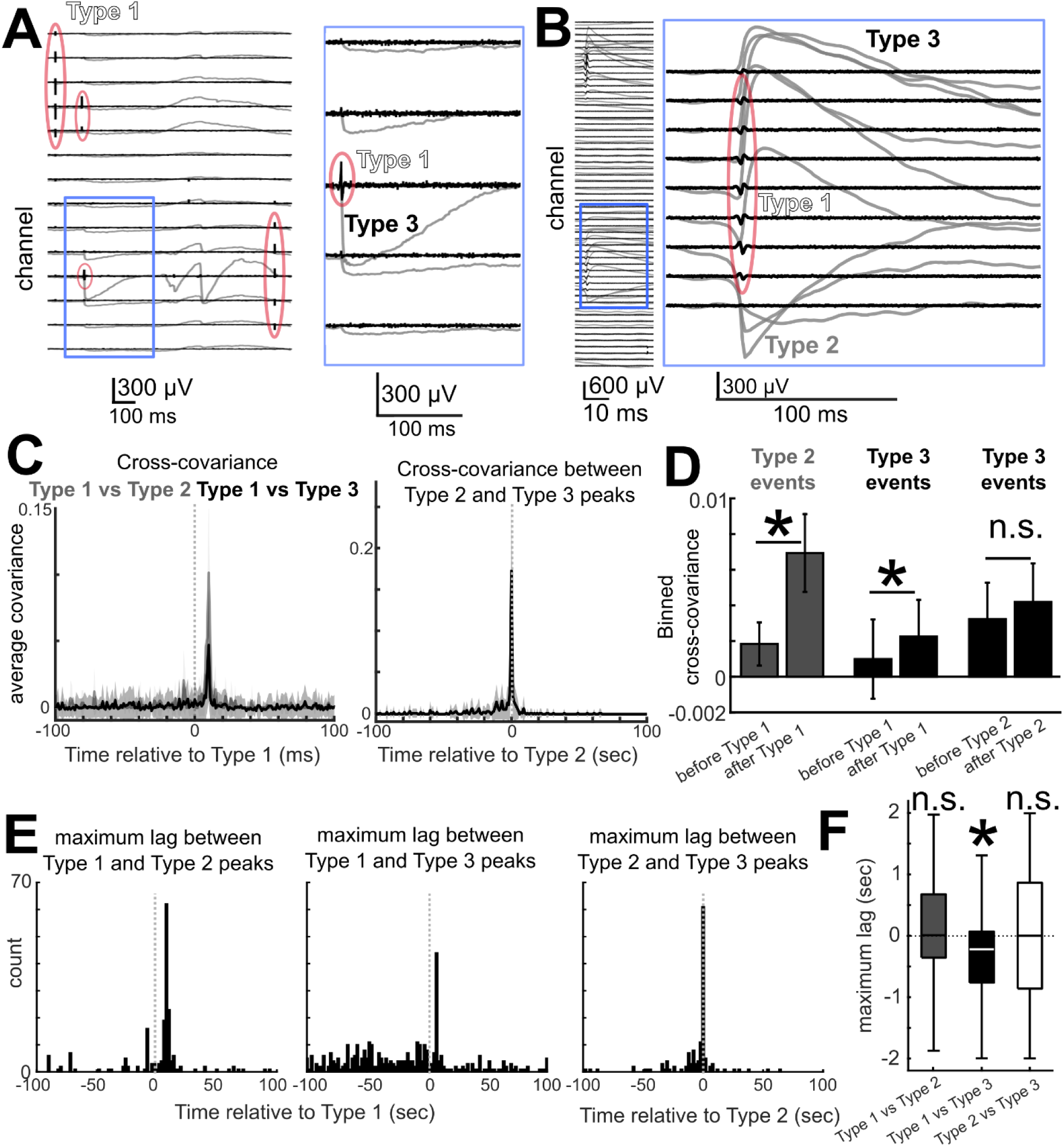
Temporal relationships between Types of events. **A-B.** Concurrent Type 1, 2, and 3 events across participants and across channels in the simultaneous high pass filtered data (between 250 and 6000 Hz, black lines) and the low pass filtered data (at 500 Hz, grey lines). Type 1 events are circled in red. Note that A shows an example in which there are Type 1 events isolated from Type 2 and 3 and in B there is a Type 1 event on a different channel from the Type 2 event. **C.** Left: Average cross-covariance plots between Type 1 and Type 2 (grey line) and Type 3 (black line) events co-occurring in time on different channels (N=24). Right: Average cross-covariance plots between Type 2 and Type 3 events co-occurring in time on different neighboring channels (N=36). Shaded lines are standard error. **D.** Average covariance from the 50 ms before each event to same interval after the event. Asterisks indicate p<0.001; N=24; Wilcoxon rank-sum test. Error bars are s.e.m. **E**. Binned maximum time lag between Type 1 event and the Type 2 and Type 3 events for all the recordings in humans (two right plots) and the binned maximum time lag between the Type 2 and Type 3 events. **F**. Median maximum lag in covariance between Type 1 and Type 2 (grey box plot), Type 1 and Type 3 (black box plot), and Type 2 and Type 3 events with confidence bounds. Asterisk indicates that distribution is significantly different from zero, p=0.0004. Wilcoxon signed rank test.

However, the detected Type 3 events spread significantly faster (Type 2: 0.05 ± 0.038 m/sec; Type 3: 0.08 ± 0.0511 m/sec; p=0.005; rank-sum=405; Wilcoxon rank-sum test) **Fig. 4Biv**). The spatial spread differences between the larger circular grid and the smaller 2-column grid (as indicated by the white versus black scatter plots; **Fig. 4Bii**, **4iii**) likely resulted in the considerably varied event speeds per participant (**Fig. 4Biv**).

### Types 2 and 3 events are detectable with other microelectrode types and near the cortical surface

To determine if these events are only present in PEDOT:PSS recordings from the cortical surface, we applied our template matching to concurrent clinical electrodes (3-mm diameter contacts on the surface of the brain; N=9). We found almost no Type 2 (n=12; 0.0027 ± 0.0061 events per second) or Type 3 (n=75; 0.0167 ± 0.0287 events per second) events on the clinical electrodes across all cases, which was significantly fewer than in concurrent PEDOT:PSS recordings (Type 2: n=1,383; 0.31 ± 0.34 events per second; Type 3: n=12,182; 2.71 ± 3.28 events per second; p<0.002 for both Type 2 and 3; Wilcoxon rank-sum test; **Fig. 4Ci**).

While the Type 2 and Type 3 events were not easily observed with clinical electrodes, we tested whether we could observe these events with other neurorecording systems. To this end, we performed the same template matching on neural recordings from patients with epilepsy semi-chronically implanted with Utah arrays, which were either 1 mm deep (N=4) or 1.5 mm deep (N=4; **Fig. 4Cii**; **Supplementary Table 3**). We found more Type 2 and Type 3 events with the shorter Utah electrodes (Type 2: n=484; Type 3: n=31,678) than the deeper ones (Type 2: n=105; Type 3: n=1,743) using the same template matching as that employed in the PEDOT:PSS data analysis (Type 2: p = 0.0209; Type 3: p = 0.0833; Type 2: 1.0 mm- 0.0640 ± 0.0391 versus 1.5 mm- 0.0123 ± 0.0083 per second; Type 3: 1.0 mm- 4.73 ± 5.6867 versus 1.5 mm- 0.18 ± 0.2678 events per second; p<0.001; N=4 each Utah array depth; Wilcoxon rank-sum test; **Fig. 4Cii**). We also used the same template-matching approach on semi-chronic laminar microelectrode data which sample multiple cortical layers (Ulbert et al., 2001; N= 9; **Fig. 4Ciii**). We observed few Type 2 events in the laminar recordings (n= 24, 0.002 ± 0.002 events per second), but we did detect more Type 3 events (n=3,084, 0.26 ± 0.46 events per second). Interestingly, the laminar recordings yielded more Type 3 events in the superficial cortical layers as was evidenced by the average negative correlation between channel numbers and Type 3 counts across the array (rho= −0.48 ± 0.38; significantly different to zero; p = 0.0313; Wilcoxon signed rank test; N= 9; **Fig. 4**). To summarize, Type 2 and Type 3 events were detectable by other than PEDOT:PSS microelectrodes, including semi-chronic electrodes, and occurred more often in top cortical layers.

### Spectral Dynamics Surrounding Type 1, 2 and 3 Events

We performed four separate types of analysis to specifically determine if any of these events were part of an ongoing oscillation (Cole and Voytek 2017). We examined 1) event-locked spectral power increases surrounding the events (performed for Type 1, 2, and 3 events); 2) peaks in power for oscillations at specific frequency bands surrounding each event (after removing the events, performed for Type 2 and 3 events, including with shuffled trials); 3) phase-locking of Type 2 and 3 events with specific frequency bands (after removing the events, performed for Type 2 and 3 events); and 4) periodic regularity in the IEI and autocorrelation plots (performed for Type 1, 2, and 3 events). We focused on recordings of spontaneous activity without additional stimuli or manipulations and on five different frequency bands (2-4 Hz, 4-8 Hz, 8-15 Hz, 15-30 Hz, 30-55 Hz, 65-200 Hz). We discuss the results of the analyses for event-locked spectral power increases. We examined power changes surrounding events, the phase-locking measures, and periodic regularity in the IEI dynamics in the **Supplementary Material** (**Supplementary Fig. 13-17**). In all the spectral analyses, we could not find a there was no significant relationship in phase or power between ongoing oscillations and Type 2 and 3 events.

All three events types showed a broadband increase in power during the event or slightly beforehand (on the order of 0.25 sec) in the event-triggered normalized power dynamics (**Supplementary Fig. 13**). Type 1 events coincided with an average broadband increase in power for the duration of the event for multiple frequency bands (4-30 Hz) (**Supplementary Fig. 13**). Type 2 events coincided with a significant broadband increase in power slightly before and during the event (p<0.001; Wilcoxon rank-sum test; corrected for multiple comparisons with false discovery rate control; **Supplementary Fig. 13**). Type 3 event-triggered LFP power had low-frequency power peaks surrounding each event, but within 0.25 sec, which were smaller than the change in power during the Type 2 events (**Supplementary Fig. 13**). Outside the 0.25-sec surrounding window, we could not find any consistent modulation of power in any frequency band preceding the events in the event-triggered power dynamics. We wanted to focus on relationships between ongoing oscillations relative to the Type 2 and 3 events and we found no clear relationship between these events and oscillations. We did not perform any tests of the spectral relationships between Type 2 and 3 events relative to external stimuli as we were attempting to study Type 1-3 events relative to ongoing oscillations that are not stimulus-linked. More importantly, the relationships of the Type 2-3 events to spectral dynamics are outside the scope of the study and will be a part of future studies, particularly in other species.

A confound, however, with these results, is that these significant changes in power could just be an effect of the sharp deflection at the onset of the Type 1, 2, and 3 events having a broadband increase in power in all frequencies. To test this, we performed the same event-triggered power calculation on the few artificial ‘events’ in saline recordings which do not have as sharp waveform onsets (**Materials and Methods**) and found no such power increase at the event onset (**Supplementary Fig. 14**).

### Type 1-3 events are not a purely human phenomenon and are present in healthy cortex

To determine if these events do not represent normal neural activity as they could be small-field inter-ictal discharges (IIDs) in epileptiform tissue (Curtis et al. 2012), and to prepare for future, more extensive mechanistic investigations (Barry 2015; Moore et al. 2017; Suzuki and Larkum 2017; Abdelfattah et al. 2019), we searched for these events in animal recordings with PEDOT:PSS electrode arrays. These studies included recordings from the barrel cortex region of the primary somatosensory cortex (S1) in alpha-chloralose anesthetized mice (N=4), the primary visual cortex (V1) in a ketamine-anesthetized mouse, the primary visual cortex (V1) in two isoflurane-anesthetized mice, and both primary and higher visual areas (VI and V4) in an anesthetized non-human primate (NHP) in an acute setting (N=1) using the circular, 2-column, and square grid PEDOT:PSS electrodes (**Supplementary Fig. 4, 5, 18**; **Supplementary Table 4**). In both the NHP and mouse barrel cortex where the array way placed directly on the cortical surface (with the dura removed), we found Type 1, 2, and 3 waveforms across recording sessions performed with different electrodes as well as repeatedly in the same recording sessions. In the V1 mouse epidural recording session where the array was placed on top of the dura, we found only Type 2 and 3 waveforms, possibly due to the low-pass filtering property of the dura (**Supplementary Fig. 18;** Type 1 activity data not shown). The Type 2 and 3 all had waveform characteristics (τ_rise time_ and τ_fall time_) and frequencies similar to those observed in the human recordings (**Supplementary Fig. 18-20**). We also performed event-triggered spectral comparison and found dynamics similar to those in humans (**Supplementary Fig. 20**). As a final test, we performed postmortem recordings in the V1 area of cortex in a mouse that was earlier anesthetized with ketamine. We found the Type 1, Type 2, and Type 3 waveforms had completely disappeared after euthanasia supporting their physiological origin (**Supplementary Fig. 18**).

### Auditory stimulation increases Type 3 events

To test if these events were relevant to neural processing during behavior, we recorded neural activity over the human superior temporal gyrus (STG) and lateral prefrontal cortex while presenting regular auditory cues in a subset of participants undergoing awake resective surgeries (Ganji et al., 2017; Hermiz et al., 2018) (N=11; **Fig. 5A**). We detected both evoked potentials and Type 1-3 events related to the auditory task (**Fig. 5B**; see **Materials and Methods**). When we performed the peri-stimulus time histogram (PSTH) calculations for Type 1-3 events, we found the Type 3 events were more frequent following the auditory stimulus (**Fig. 5B**). We found no significant difference in counts between Type 1 and Type 2 events (**Fig. 5B**, event rate (Hz) comparison: p=0.1078; Wilcoxon rank-sum test). Further behavioral and auditory stimuli may be needed to further distinguish these dynamics, particularly since spectral analyses have shown there are significant changes in high gamma and other frequency bands in response to auditory stimulation over the STG intraoperatively using PEDOT:PSS (Hermiz et al. 2018). Additionally, waveform characteristics did not vary significantly during the auditory cue versus outside the cue epochs (**Supplementary Fig. 21**).

### Pharmacological manipulation alters event rate

We hypothesized that if these events are physiological, then Type 1, 2, and 3 events should vary in rate with cold saline application (N=8) or intravenously administered pro-convulsant agents (methohexital or alfentanil), which are used to map seizure foci in the brain (Curtis et al. 2012) (N= 9; **Supplementary Table 1-2**). Across participants, both cold saline and pro-convulsant medication such as methohexital suppressed event frequency (**Fig. 5C-D**).

Administering pro-convulsant medication notably lowered the rate of Type 2 and 3 events while still increasing epileptiform activity. More specifically, Type 2 events tended to decrease four minutes after the addition of pro-convulsant medication (0.0.0096 ± 0.0772 Hz compared to 0.02 ± 0.219 Hz before) (p=0.033; N=9; Kruskal-Wallis multiple comparisons test). Type 3 events were significantly less frequent three minutes after pro-convulsant medication was intravenously injected (0.52 ± 0.813 Hz before injection vs 0.38 ± 0.678 Hz four minutes after injection and continuing up to 5 minutes; N=9; p<0.0001; Kruskal-Wallis multiple comparisons test; **Fig. 5C-D** **, left column**). Type 1 events were only found in 1 of the 9 recordings with pro-convulsant medication, so we could not perform the same statistical comparisons.

We contrasted these results with changes in the rate of interictal discharges (IIDs) as identified by trained epileptologists. IIDs were significantly more frequent after intravenous injection of pro-convulsant medication (0.001 ± 0.001 Hz before injection vs 0.0099 ± 0.0094 Hz three minutes after injection; N=9; **Fig**. **5C-D, left column**), adding further evidence that these events belong to healthy electrophysiological repertoire rather than miniature IIDs (Schevon et al. 2008, 2010).

Adding cold saline suppressed Type 1, 2 and 3 events across participants (N=8; **Fig. 5C-D**). Type 1 event counts dropped significantly, from 0.0029 ± 0.0023 Hz to 0.0001 ± 0.0002 Hz two minutes after cold saline application (p<0.0001; Kruskal-Wallis multiple comparisons test). Type 2 event rate decreased from 0.01 ± 0.024 Hz during baseline to 0.002 ± 0.003 Hz two minutes after cold saline application across all channels across participants, though the change was not significant (p=0.7282; Kruskal-Wallis multiple comparisons test). Cold saline also rendered Type 3 events less frequent, with a significant decrease after two minutes (0.04 ± 0.046 Hz before addition of cold saline vs 0.01 ± 0.019 Hz two minutes after saline addition; p<0.0001; Kruskal-Wallis multiple comparisons test; **Fig. 5C-D**, right column).

It should be noted that neither intervention significantly altered waveform characteristics relative to baseline (p>0.12; Wilcoxon rank sum test; **Supplementary Fig. 21**). Additionally, in all cases, room temperature saline was regularly used to irrigate the cortical surface during the surgery. There was no change in event rate during application of room temperature saline (data not shown).

### Electrical stimulation changes the rates and spread of events

To further test the hypothesis that Type 1-3 events are neural in origin, we examined changes in activity during and after trains of electrical stimuli (as in motor or language mapping (Berger and Ojemann 1992; Mueller and Morris 1993; Kawaguchi et al. 1996; Viventi et al. 2011; Borchers et al. 2012), **Fig. 5E**). When trains of stimuli were applied to the surface of the cortex, induced travelling waves of Type 3 events were observed progressing across all channels, sometimes starting soon after the end of stimulation but often initiating 1 to 2 seconds after the stimulation (**Fig. 5E**). These propagating waves of activity were observed in all instances with clinically relevant trains of electrical stimulation. Similar activity waves were reported in other situations in other cortical recordings (Viventi et al. 2011), though not due to stimulation.

We examined the PSTH of Type 1-3 events around each stimulation train (normalized to the 1 second before stimulation) and found that, during stimulation, Type 2 event rate decreased and Type 3 events were significantly inhibited (p<0.0001; Wilcoxon rank-sum comparison between baseline and each binned time window; **Fig. 5F-G**). Immediately after stimulation, Type1, 2 and 3 events were all more frequent (p<0.0001; Wilcoxon rank-sum comparison between baseline and each binned time window; **Fig. 5F-G**) while waveform characteristics were unchanged (**Supplementary Fig. 22**).

### Temporal interactions between different event types

We examined whether the Type 1 events had a consistent relationship to the slower Type 2 and 3 events (**Fig. 6**). We found several examples across participants where either Type 1, 2, and 3 events overlapped in time or Type 1 events preceded Type 2 or 3 events (**Fig. 6A-B**). Further, we found small, but prominent, peaks in event cross-covariances with Type 1 events preceding Type 2 and Type 3 events, often by ∼10 ms (**Fig. 6C, D**; p<0.0015; Wilcoxon rank-sum test; N=36). The small cross-covariance peak was present between Type 2 and Type 3 events, though there was no significant lag between the two (**Fig. 6C, D**; p=0.25). Another way to measure the timing relationship between the events was to identify the maximum time lag between Type 1 events and Type 2 and Type 3 events for each recording and determine the count and distribution of the maximum lag. Again, we found a delayed peak around 10 ms between Type 1 and both Type 2 and 3 events (**Fig. 6E**). Interestingly, however, there were also a number of Type 3 events preceding Type 1 events at longer time scales (closer to 50-100 ms; p<0.0001; Wilcoxon signed rank test for significant different to zero; N=24; **Fig. 6F**). We also found a significant deflection in the LFP around Type 1 events (after normalizing relative to 0.5 sec before the fast event) across participants (p<0.05, false discovery rate controlled; N=24; **Supplementary Fig. 22**). Together, these findings suggest that Type 1 events generally occur prior to Type 2 and 3 events at the ∼10 millisecond time scale.

## Discussion

We observed repeated, localized microscale waveforms across both surface and penetrating microelectrode recordings in human and non-human cortex, and these waveforms could be divided into faster (∼1 ms, Type 1) and slower (∼10-100 ms, Type 2 and Type 3) events. To exclude possible artifacts of the recording system, electrodes, and other confounding factors, we examined recordings in eight different centers, across 54 patients, in four mice and an NHP, and with three different recording systems. We found these events in all examined cases irrespective of the type of recording microelectrodes (PEDOT:PSS, Utah, and laminar electrodes) and under both acute and semi-chronic conditions. The different event types were selectively and separately inhibited by pro-convulsant medication and cold saline and were differentially impacted by electrical stimulation and sensory stimuli. In addition, we detected the Type 2 and 3 events more often in the upper cortical layers of the cortex, whether with shorter Utah arrays or in the superficial contacts of laminar microelectrode arrays. Such events were essentially absent in non-biological recordings (e.g. in saline and after euthanasia or in “upside down” recordings), or in concurrent neural recordings using 3 mm diameter clinical macroelectrode contacts. Furthermore, these events had characteristic spatiotemporal patterns, were repeated and localized, and showed rare but statistically significant interactions, with Type 1 events more often leading Type 2 and 3 events.

Given that recordings in humans were performed during clinically necessary intracranial monitoring, a significant challenge for interpreting these results is the question of whether the Type 1, 2, and 3 events reflect normal brain activity or only represent pathology. If these events were indeed related to abnormal brain activity, they would still be of interest as biomarkers of pathology; however, the present results provide substantial evidence that these are physiologically normal events. First, pathological events are unlikely to be induced by auditory stimuli. Second, we found these events in all examined patients regardless of the reason for their surgery or the location of recording as well as during semi-chronic recordings using Utah arrays and laminar recordings. Third, we observed similar events in both rodents and an NHP. Taken together, these findings strongly suggest that these events, recorded from the cortical surface, could be found in normal tissue and represent normal physiological activity.

Importantly, these surface events appeared as stand-alone occurrences with asymmetric waveforms and relatively fast kinetics, but without rhythmicity, indicating they were not part of typical ongoing oscillations. Were events part of an ongoing oscillation, we would expect a longer-term increase in power before and after the event, either locked to the event onset or related to the phase of an ongoing oscillation. Yet, no significant relationships were observed between ongoing oscillations and these different events in either power or phase relationships (background oscillations were observed separate from these discrete events). This supports the notion that these events occurred independently rather than as fragments of ongoing oscillations. Conversely, this is not to say that the events might not have some relationship to oscillatory activity under certain circumstances but, like in the traditional analysis of single unit activity, determining such a relationship would require specialized analysis focusing on a particular oscillation frequency band in a specific brain state or context. Furthermore, we note that more typical oscillations were observed in these recordings and again, these were independent of the unitary events described here.

One caveat of the present work, however, is that we are likely under-sampling Type 2 and 3 events through our strict set of criteria and template matching. Thus, our interpretation of the rates and spatial characteristics of the different events might not be fully accurate. As has occurred with spike sorting, future work will necessarily involve an evolution toward ever more sophisticated tools to detect these events in the course of spontaneous neural activity. An additional related potential confound is that the majority of the data set is from acute recordings in the OR; an experimental arena prone to considerable noise and instability for electrophysiological recording. To mitigate this issue, we repeated these tests across hospitals, participants, species, electrodes, and recording systems. Even then, concerned that the environment could be introducing noise, we examined semi-chronic data to demonstrate the presence of these events. However, even with all these tests, we concede that some noise could be detected as events. Again, further, more sophisticated detection approaches could clear up these issues especially as we obtain a greater understanding of the cellular mechanisms underlying this activity. We do hypothesize, however, that the low impedance conformable spatially high resolution PEDOT:PSS electrodes are better able to spatially map these events than rigid devices on the cortical surface, but this will require further testing.

Of course, the cellular processes underlying these events are unknown and warrant further investigation. Yet, it is worth speculating on several possibilities. One is that the events reflect activity in specific, specialized, neural cell types present in supragranular layers of the cortex (Rakic and Zecevic 2003; Zaitsev et al. 2009; Mohan et al. 2015; Gabbott 2016; Boldog et al. 2018). This could include Cajal-Retzius cells or other relatively rare cell types whose cell bodies are found in upper cortical layers (Rakic and Zecevic 2003; Mohan et al. 2015; Gabbott 2016). Alternatively, these events could reflect ionic flux in glial cells in upper cortical layers – although the fast kinetics make this somewhat less likely as astrocytic waves normally occur on the time scale of seconds (Hassinger et al. 1996; Scemes and Giaume 2006; Takata and Hirase 2008; Kuga et al. 2011; Khakh and McCarthy 2015; Rouach et al. 2018).

More intriguing is the possibility that some of the Type 1 and faster Type 2 events reflect back-propagating dendritic action potentials (Robbins et al. 2013; Stuart and Spruston 2015; Moore et al. 2017; Jia et al. 2019) while the slower Type 2 and Type 3 events could be the result of dendritic calcium spikes (Bakkum et al. 2013; Barry 2015; Suzuki and Larkum 2017). A similar type of activity has been reported using fine electrodes in freely behaving rats. In that instance, putative dendritic action potentials were recorded using tetrodes following the formation of a glial sheath to increase the contact between neural dendrites and the fine wires (Moore et al. 2017). The waveform shapes of extracellular dendritic action potentials reported in that study had similar voltage range (∼.5-1 mV) to the Type 2 events described here, although their duration was shorter (duration ∼5-10 ms). Type 2 and/or Type 3 events could also be indicative of localized calcium spikes in the apical tufts of pyramidal neurons from deeper layers (Ross 2015; Suzuki and Larkum 2017; Golding et al. 2018). Similar time courses and waveforms have also been recorded from the cortex surface in anesthetized rats, phenomena that were then verified, through optogenetics, laminar probes, pharmacological manipulation, and other tools including calcium imaging, as dendritic calcium spikes from layer 2/3 or layer 5 pyramidal neurons (Suzuki and Larkum 2017). Extrapolating from those findings and the data presented here, it is possible that high density microelectrodes could be recording dendritic activity at the pial surface.

In addition, some of the Type 1 waveforms were within the range of extracellular action potential dynamics. However, Type 1 events also included very narrow waveforms, some with a triphasic appearance and some with relatively large amplitudes compared to previously recorded somatic action potentials. It should be noted that primate pyramidal neurons have been shown to produce fast (or “thin”) spikes not often observed in non-primate species (Vigneswaran et al. 2011). The different waveform shapes for the fast events we identified may also represent different neural subclasses (Bartho et al. 2004; Vigneswaran et al. 2011; Chan et al. 2014; Trainito et al. 2019). In addition, with their short duration and triphasic morphology, some Type 1 events resemble axonal action potential recordings (Robbins et al. 2013; Stuart and Spruston 2015; Jun et al. 2017). In this scenario, some of the Type 1 events would reflect activity in unmyelinated axons in the uppermost cortical layers (Mohan et al. 2015). This notion is supported in part by the observed relationship whereby Type 1 events preceded some of the Type 2 and 3 events, thereby implying that the fast Type 1 events are axon inputs while the slower events could be a post-synaptic consequence. These possibilities may also account for the fact that some of the Type 1 and Type 2 and 3 events were spatially co-localized. While this colocalization was rare across recordings, the timing relationships were significant.

In summary then, we speculate that these different classes of relatively fast activity could reflect a range of neuronal events including axonal action potentials, fast and slow events in the dendrite and somatic action potentials. In the future, optical calcium and voltage imaging in combination with electrophysiological recording approaches in animal models will be essential to further resolve the mechanisms of generation of these events (Harvey et al. 2009; Roome and Kuhn 2018; Abdelfattah et al. 2019; Adam et al. 2019; Chiang et al. 2020; Hossain et al. 2020)(1, 3, 63–67). Electrophysiological events will need to be examined in awake animals using calcium imaging (Suzuki and Larkum 2017), intracellular recordings (Harvey et al. 2009; Abdelfattah et al. 2019), pharmacological interventions (Suzuki and Larkum 2017), and genetic cell-type-specific manipulation (Abdelfattah et al. 2019).

Examination of the entire neurophysiological signal from microelectrode technologies, can uncover, or at least highlight, signals otherwise understudied in the neuroscience field. These results open the possibility that we can access circuit-level dynamics in the cortex using high resolution sampling. If so, then we could more thoroughly study the detailed neural circuit activity underlying physiological events and build more complete bridges from the single cell level to large scale population behavior in the context of both healthy and pathological human brain dynamics.

## Acknowledgments

We would like to thank Yangling Chou, Erica Johnson, Melissa Murphy, Aaron Tripp, Fausto Minidio, Alex Zhang, and Gavin Belok for help in data collection and, of course, the patients for their willingness to participate in this research. This research was sponsored by the U.S. Army Research Office and Defense Advanced Research Projects Agency under Cooperative Agreement Number W911NF-14-2-0045. In addition, other support included National Institutes of Health under Award Number 1F32MH120886 to DRC, ECOR and K24-NS088568 to SSC, Tiny Blue Dot Foundation to SSC and ACP, and by an NSF-CAREER award #1351980, NSF CMMI award #1728497, and NSF-ECCS EAGER award #1743694 and an NIH award # DP2-EB029757 to SAD, and BRAIN Initiative R01MH111359 to AD, and NIH (NEI R01-EY029022 / EY023651 and NINDS U01-NS099700) and by the Dept. of Defense / CDMRP (VR170089) to SBR, SWL, and SIF. Other support includes the Hungarian Brain Research Program: 2017-1.2.1-NKP-2017-00002 to IU. The views and conclusions contained in this document are those of the authors and do not represent the official policies, either expressed or implied, of the funding sources.

## Author Contributions

Conceptualization, A.C.P., J.C.Y., S.S.C., S.A.D. Methodology, J.H., M.G., A.C.P., J.C.Y., S.S.C., S.A.D. Formal analysis, A.C.P., J.C.Y., D.J.S., M. H., Investigation, A.C.P., J.C.Y., D.R.C., N.R., K.K., S.B.R., S.W.L., D.M., S.I.F., P.S.J., B.V.N., S.B-H., A.M.T.R., D.S., D.P.C., I. U., D. F., O. D., J. R. M., D. L. S., E. N. E., J. L., S. K. B., Z.M.W., G.R.C., S.A.D., S.S.C. Resources, S.A.D., S.S.C., S.I.F, S.W.L., S.H.L., M.G., Y.G.R., H.O., L.H., J. L., Y. T. Data curation, A.C.P., D.J.S., J.C.Y., A. R. O. Writing, A.C.P., J.C.Y., S.S.C., S.A.D., D.P.C., Z.M.W., S.B.R., N.R., A.D., D.J.S., D.R.C., J.L., G.R.C., S. I. F., S. K. B., A. R. O. Visualization, A.C.P., D.J.S., M. H. Supervision, S.A.D., S.S.C. Project administration, A.C.P., J.C.Y., D.R.C., S.A.D., S.S.C. Funding acquisition, S.A.D., S.S.C., S.I.F., S.W.L., I. U.

