## Supplemental Material for "Microscale physiological events on the human cortical surface"

**Supplemental material included:**

**Supplementary Material for validation of Type 2 and 3 events: Recording using a different system and over a putative tumor**

**Supplementary Material for examination of the relationship between oscillations and Type 1-3 events**

**Supplementary Video 1. Example LFP dynamics shown on a PEDOT:PSS grid over time.** Recordings per good channel shown on the left and the same recordings mapped to the circular grid on the right. Distance between lines in the leftmost grid: 200 µVolts. The color coding of the dots of the channels shown in the left grid (along the y axis) matches the small single point (dot) channel color coding underlying the traces on the right. Data are low pass filtered at 500 Hz and sampled at 1000 Hz.

**Supplementary Table 1. Distribution and numbers of cases and treatments**.

**Supplementary Table 2. Information per participant and recording. 2-column and circular refer to PEDOT:PSS grid configurations.**

**Supplementary Table 3. Information per participant and recording, Utah array and laminar recordings.**

**Supplementary Table 4. Information per mouse and NHP recording.**

**Supplementary Figure 1. Right hemisphere 2-column laminar 50 µm pitch PEDOT:PSS electrode locations**.

**Supplementary Figure 2. Left hemisphere 2-column laminar 50 µm pitch PEDOT:PSS electrode locations**.

**Supplementary Figure 3. PEDOT:PSS electrode locations of the 800 µm pitch 2-column laminar and the circular grid electrodes.**

**Supplementary Figure 4. Example RMS examination of channel activity during the quiet periods of a recording and impedance changes of the PEDOT:PSSelectrodes during each conditional step.**

**Supplementary Figure 5. Locations of electrode channels per electrode which survived quality control checks, 2-column grid.**

**Supplementary Figure 6. Locations of electrode channels per electrode which survived quality control checks, Circular and rectangular grids**.

**Supplementary Figure 7. Inter-Event-Interval (IEI) distribution for the Type 1 events.**

**Supplementary Figure 8. Principal components analysis (PCA) on voltage waveforms to determine if there were different clusters of waveforms.**

**Supplementary Figure 9. Criteria for detecting Type 2 and 3 events and variation across channels versus within channels.**

**Supplementary Figure 10. Detecting and separating Type 2 and Type 3 waveforms in saline recordings versus on-brain recordings using the same electrodes in the operating room**.

**Supplementary Figure 11. Type 2 and Type 3 waveforms in saline recordings versus on-brain recordings using the same electrodes in the operating room**.

**Supplementary Figure 12. Saline recording waveform metrics compared to recordings on brain with the same sets of electrodes**.

**Supplementary Figure 13. Event-triggered spectral changes in the local field potential (LFP)**.

**Supplementary Figure 14. Event-triggered time-locked power dynamics, saline.**

**Supplementary Figure 15. Power in different frequency bands surrounding Type 2 and 3 events.**

**Supplementary Figure 16. Phase-locking to different frequency bands per event type per frequency band**.

**Supplementary Figure 17. Inter-Event-Interval (IEI) and autocorrelation dynamics for the Type 2 and Type 3 events.**

**Supplementary Figure 18. NHP and mouse recordings.**

**Supplementary Figure 19. NHP and mouse recordings, waveform characteristics.**

**Supplementary Figure 20. Event-triggered time-locked power dynamics for Type 2 and 3 events across NHP and mouse recordings**.

**Supplementary Figure 21. Waveform measurements during the application of pro-convulsant medication, cold saline, auditory stimulation and electrical stimulation.**

**Supplementary Figure 22. Spike-triggered voltage averages around Type 1 events.**

***Supplemental Material for validation of Type 2 and 3 events: Recording using a different system and over a putative tumor***

Additional evidence for the idea that these Type 2 and 3 events were neural in origin by first recording from normal brain tissue, and then, in the same session, recording from what appeared to be a tumor. We found Type 2 (n=2) and 3 (n=187) events when recording over the normal brain tissue. After moving the electrode over the tumor, we found no Type 2 events and found a significant decrease in Type 3 frequency and amplitude as well as a significant increase in both the time to rise and time to fall of the Type 3 events over tumor tissue ($p<0.05$; Wilcoxon rank-sum test). In other words, over tumor tissue, we were far less likely to detect Type 2 and 3 events, and when detected, their waveforms were both smaller and significantly slower than those recorded over normal brain tissue.

To verify that these events were not novel artifacts created by our custom-fabricated, Intan-based recording system, we performed simultaneous recordings using two circular grid electrodes, one recording using the Intan system and the other recording using a Blackrock NSP, a commonly used neurophysiological system (two circular electrodes in **Supplementary Fig. 3**). We did indeed find both Type 2 and 3 events while recording simultaneously on the same brain, which confirms that they are not artifacts of the recording system (data not shown), indicating that we were not simply recording events (data not shown).

Finally, we performed an upside-down recording (N=1), where we recorded from cortex with the PEDOT:PSS electrode recording neural tissue and then flipped the recording upside down on the surface of the cortex. The idea was to determine if the added noise in the operating room could be introducing the Type 2 and 3 events. The number of detected events with the right side up recording: n=2 Type 2 and n=446 Type 3 events with the upside down recording had no Type 2 events and only 48 Type 3 events for the same 5 minute duration.

***Supplemental Material for examination of the relationship between oscillations and Type 1-3 events***

To address this broadband effect of the waveform onset, and as an additional test of the spectral relationships between ongoing oscillatory activity and Type 2 and 3 events, we calculated power for 10-second windows of time but excluded the 0.2 sec before and after the onset of the event.  We also performed the same power analysis on random, shuffled times, shifting the 'onset time' ±10 sec per event.  This approach allowed us to evaluate whether there were consistent power increases in oscillations surrounding each event. The power curves did not vary significantly different between these different conditions across cases and frequencies nor did the adjusted time window (moving every 0.1 sec) power spectra differ for the shuffled versus the event-locked power dynamics. Indeed, there were essentially no significant differences between shuffled versus event-locked power spectral analyses ($p<0.001$; corrected for multiple comparisons; **Supplementary Fig. 15**). Therefore, beyond the initial sharp change in voltage at the onset of the Type 2 and 3 events, we could not find any consistent peaks or decreases in power at any frequency band.

Next, we examined Type 2 and 3 events phase-locking to oscillations in specific frequency bands. After performing a Hilbert transform per frequency band, we calculated the average phase-locked value (vector) per event Type per channel. Mean vector lengths for the phase values were low (far below a mean vector of 1 that would signify a strong phase relationship; **Supplementary Fig. 16**) and did not have significant unipolar distributions ($p>0.05$, Rayleigh test, $N=36$). In other words, there was no significant phase relationship between ongoing oscillations and Type 2 and 3 events. Finally, we examined the periodicity of the IEIs and auto-correlations and found no consistent periodicity for any event type (**Supplementary Fig. 17**).

**Supplementary Table 1**. **Distribution and numbers of cases and manipulations**.

| Case type with concurrent PEDOT:PSS recording | Number |
|---|---|
| Not-awake cases | 22 |
| Awake cases | 15 |
| Auditory or cognitive task performed | 12 |
| PEDOT:PSS strip electrodes tested | 26 |
| PEDOT:PSS circular grid electrodes tested | 11 |
| Epileptiform-inducing medication applied | 9 |
| Cold saline applied | 8 |
| Number of cases with clinical electrical stimulation | 9 |
| Number of cases with research electrical stimulation | 4 |
| Number of cases not analyzed due recording instability | 1 |
| **Total, PEDOT** | **37** |
| **Other microelectrode recordings** | **Number** |
| Semi-chronic laminar recordings, awake | 9 |
| Utah array, 1 mm depth, awake | 4 |
| Utah array, 1.5 mm depth, awake | 4 |
| **Total, Other arrays** | **17** |
| **Total participants** | ***54*** |

**Supplementary Table 2**. **Information per participant and recording. 2-column and circular refer to PEDOT:PSS grid configurations**

| ***Participant designation*** | **Case type with concurrent recording** | **Electrode type** | **Clinical recordings** | **stable recording sites** | **Duration of the recording** |
|---|---|---|---|---|---|
| *IP01* | Left lobectomy, temporal lobe | 2-column | Yes | 57/128 | Position 1: 1003<br>Position 2: 150 |
| *IP02* | Awake left craniotomy for tumor, temporal lobe | 2-column | Yes | 67/128 | Position 1: 445 |
| *IP03* | Right lobectomy, temporal lobe | 2-column | Yes | 20/128 | Position 1: 544 |
| *IP04* | Right lobectomy, temporal lobe | 2-column | Yes | 18/128 | Position 1: 1003 |
| *IP05* | Right lobectomy, temporal lobe | 2-column | Yes | 23/128 | Position 1: 628 |
| *IP06* | Right lobectomy parietotemporal | 2-column | Yes | 53/96 | Position 1: 348 |
| *IP07* | Awake left craniotomy for tumor, parietal lobe | 2-column | Yes | 93/128 | Position 1: 604<br>Position 2: 54 |
| *IP08* | Right lobectomy, temporal lobe | 2-column | Yes | 74/128 | Position 1: 225<br>Position 2: 170 |
| *IP09* | Right lobectomy temporal, parietal lobe | 2-column | Yes | 46/128 | Position 1: 870<br>Position 2: 317 |
| *IP10* | Right lobectomy, frontal lobe | 2-column | Yes | 64/128 | Position 1: 941 |
| *IP11* | Right lobectomy, temporal lobe | 2-column | Yes | 96/128 | Position 1: 301 |
| *IP12* | Awake left craniotomy for lobectomy, temporal lobe | 2-column | Yes | 89/128 | Position 1: 346 |
| *IP13* | Right lobectomy temporal, temporal lobe | 2-column | No | 36/128 | Position 1: 548<br>Position 2: 270 s |
| *IP14* | Right lobectomy temporal, temporal lobe | 2-column | Yes | 0/128 | Position 1: 1377 |
| *IP15* | Awake left craniotomy for tumor, temporal lobe | 2-column | Yes | 57/128 | Position 1: 887 |
| *IP16* | Right lobectomy temporal, temporal lobe | 2-column | Yes | 103/128 | Position 1: 1410 |
| *IP17* | Right lobectomy temporal, temporal lobe | Circular | Yes | 55/128 | Position 1: 2205 |
| *IP18* | Right craniotomy, motor strip, prefrontal lobe | Circular | Yes | 62/128 | Position 1: 300<br>Position 2: 145<br>Position 3: 146 |

| | | | | | |
|---|---|---|---|---|---|
| *IP19* | Awake right craniotomy for tumor, temporal lobe, parietal lobe | Circular | Yes | 75/128 | Position 1: 414<br>Position 2: 172<br>Position 3: 136 |
| *IP20* | Awake left craniotomy for tumor, temporal lobe | 2-column | No | 48/128 | Position 1: 445 |
| *IP21* | Awake left craniotomy for tumor, temporal lobe | 2-column | No | 41/128 | Position 1: 256 |
| *IP22* | Left craniotomy for tumor, temporal lobe | 2-column | No | 113/128 | Position 1: 176<br>Position 2: 313 |
| *IP23* | Right lobectomy temporal, temporal lobe | Circular | No | 21/64 | Position 1: 540 |
| *IP24* | Awake left craniotomy for tumor, temporal lobe | 2-column | No | 57/128 | Position 1: 380 |
| *IP25* | Right craniotomy, parietal lobe, prefrontal lobe | 2 Circular, electrodes | Yes | 99/128; 107/128 | Position 1: 1200<br>Position 2: 1500 |
| *IP26* | Awake craniotomy for tumor, temporal lobe | 2-column | No | 107/128 | Position 1: 497 |
| *IP27* | Awake craniotomy for tumor, prefrontal lobe | 2-column | Yes | 103/128 | Position 1: 1385 |
| *IP28* | Awake craniotomy for tumor, prefrontal lobe | Circular | No | 97/128 | Position 1: 762 |
| *IP29* | Awake craniotomy for tumor, temporal lobe | 2-column | No | 119/128 | Position 1: 1163.5 |
| *IP30* | Awake craniotomy for tumor, temporal lobe | 2-column | No | 60/128 | Position 1: 923.5 |
| *IP31* | Right lobectomy temporal, temporal lobe | Circular | Yes | 63/128 | Position 1: 1330 |
| *IP32* | Right lobectomy temporal, temporal lobe | Circular | Yes | 88/128 | Position 1: 945 |
| *IP33* | Left lobectomy, temporal lobe | Circular | Yes | 88/128 | Position 1: 380<br>Position 2: 920 |
| *IP34* | Right lobectomy temporal, temporal lobe | Circular | Yes | 93/128 | Position 1: 710 |
| *IP35* | Awake left craniotomy for tumor, prefrontal lobe | 2-column | No | 89/128 | Position 1: 930 |
| *IP36* | Awake left craniotomy for | 2-column | Yes | 92/128 | Position 1: 106<br>Position 2: 3200 |

| | | | | | |
|---|---|---|---|---|---|
| | lobectomy, temporal lobe | | | | |
| *IP37* | Awake left craniotomy for lobectomy, prefrontal lobe | Circular | Yes | 108/128 | Position 1: 1358<br>Position 2: 687 |

**Supplementary Table 3**. **Information per participant and recording, Utah array and laminar recordings.**

| ***Participant designation*** | **Implanted region** | **Electrode type** | **Clinical recordings** | **stable recording sites** | **Duration of the recording** |
|---|---|---|---|---|---|
| *EP01* | Right posterior superior temporal gyrus | Laminar electrode | Yes | 23/23 | Position 1: 785 s |
| *EP02* | Left medial temporal lobe | Utah array, 1.0 mm | Yes | 84/96 | Position 1: 2400 s |
| *EP03* | Left medial temporal lobe | Utah array, 1.5 mm | Yes | 95/96 | Position 1: 2400 s |
| *EP04* | Right prefrontal lobe | Utah array, 1.0 mm | Yes | 87/96 | Position 1: 2400 s |
| *EP05* | Left superior temporal lobe | Utah array, 1.5 mm | Yes | 94/96 | Position 1: 2400 s |
| *EP06* | Right medial temporal lobe | Utah array, 1.5 mm | Yes | 94/96 | Position 1: 2400 s |
| *EP07* | Left lateral prefrontal | Laminar electrode | Yes | 23/23 | Position 1: 1051 s |
| *EP08* | Left posterior cingulate | Laminar electrode | Yes | 24/24 | Position 1: 640 s |
| *EP09* | Right prefrontal lateral | Laminar electrode | Yes | 23/23 | Position 1: 1346 s |
| *EP10* | Right anterior superior temporal gyrus | Laminar electrode | Yes | 23/23 | Position 1: 2000 s |
| *EP11* | Right lateral prefrontal | Laminar electrode | Yes | 21/21 | Position 1: 2380 s |
| *EP12* | postcentral gyrus | Laminar electrode | Yes | 20/20 | Position 1: 2000 s |
| *EP13* | Left lateral prefrontal | Laminar electrode | Yes | 23/23 | Position 1: 2000 s |
| *EP14* | Right supramarginal gyrus | Laminar electrode | Yes | 23/23 | Position 1: 1200 s |
| *EP15* | Left medial temporal lobe | Utah array, 1.0 mm | Yes | 96/96 | Position 1: 2400 s |
| *EP16* | Right medial temporal lobe | Utah array, 1.0 mm | Yes | 83/96 | Position 1: 2400 s |
| *EP17* | Left prefrontal lobe | Utah array, 1.5 mm | Yes | 80/96 | Position 1: 2400 s |

**Supplementary Table 4**. **Information per NHP or mouse and recording.**

| *designation* | **Case type with concurrent PEDOT:PSSrecording** | **PEDOT:PSSelectrode type** | **stable recording sites** | **Duration of the recording** |
|---|---|---|---|---|
| *NHP01* | Isoflurane anesthetized NHP recording of visual cortex | 2-column | 64/128 | 759 s |
| *MM01* | Recording of visual cortex in an mouse anesthetized with ketamine | Circular | 77/128 | 1465 s |
| *MM01Euth* | Recording of visual cortex in a euthanized mouse | Circular | 89/128 | 1486 s |
| *MM02* | Alpha-chloralose anesthetized mouse recording of mouse barrel cortex | Square | 29/32 | 3236 s |
| *MM03* | Recording of visual cortex in a mouse anesthetized with isoflurane | Circular | 80/128 | 270 s |
| *MM04* | Recording of visual cortex in a mouse anesthetized with isoflurane | Circular | 105/128 | 290 s |

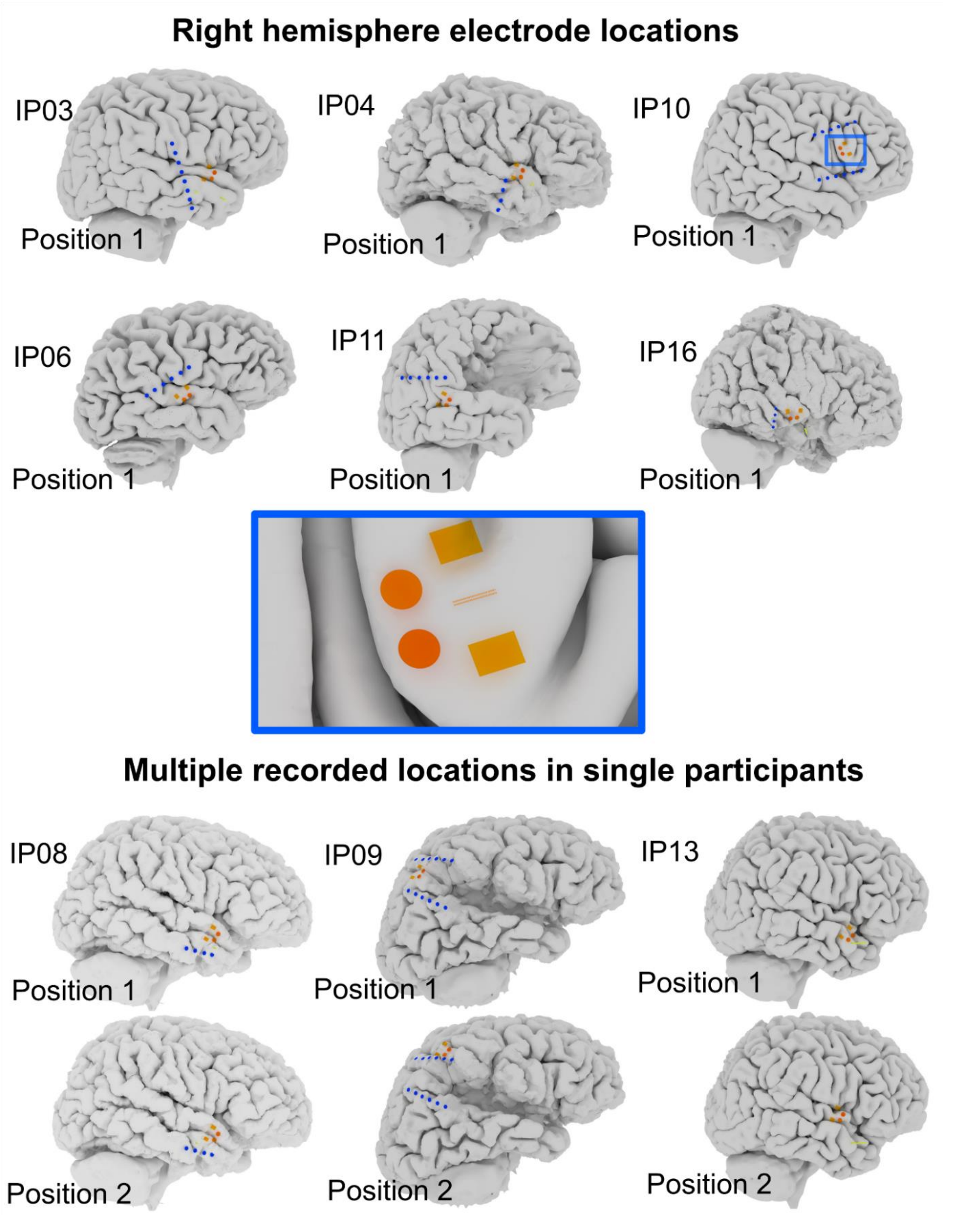

**Supplementary Fig. 1. Right hemisphere 2-column laminar 50 µm pitch PEDOT:PSS electrode locations**. Orange electrode models indicated on the brain across participants. Blue electrodes are clinical leads.

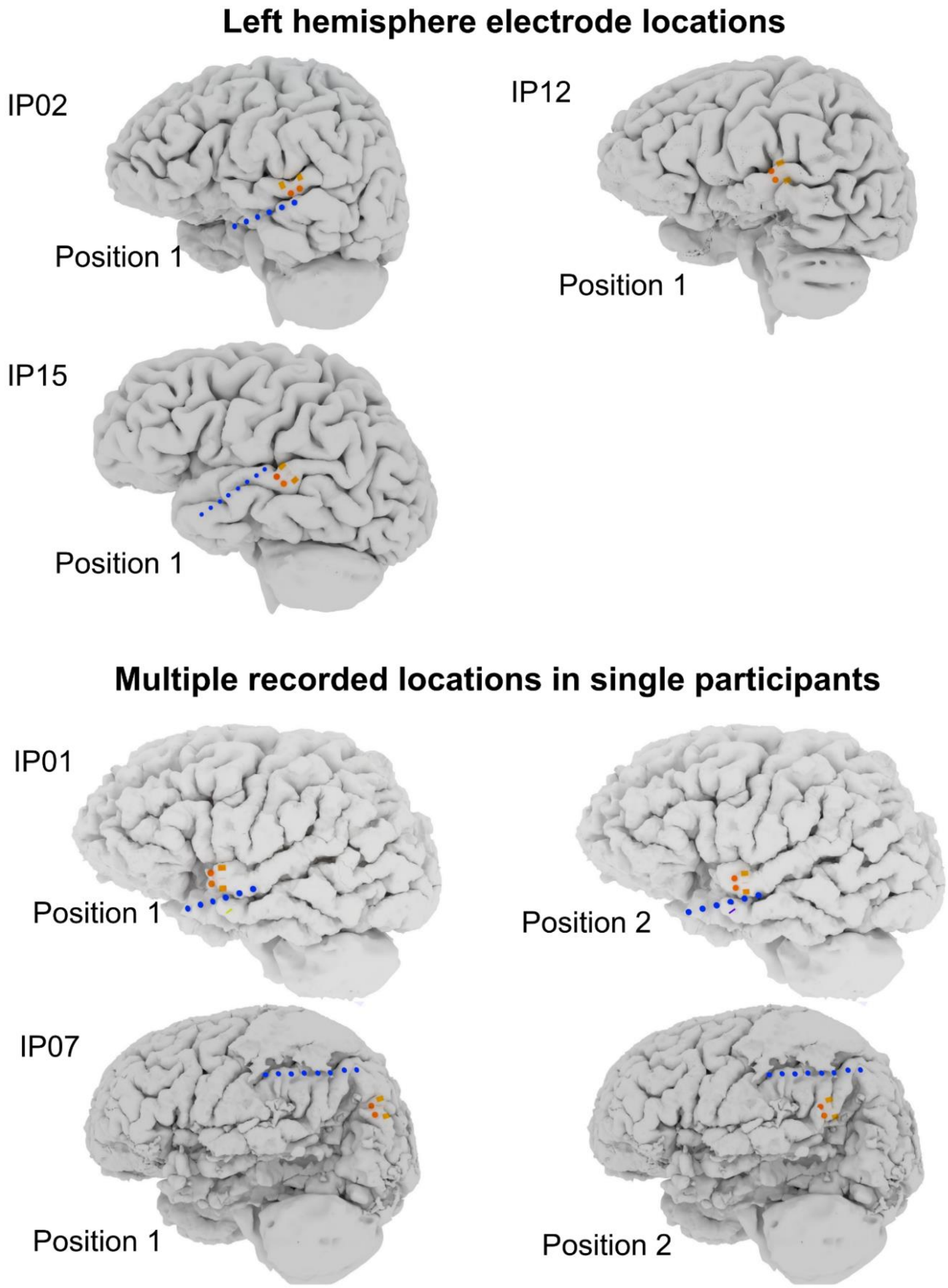

**Supplementary Fig. 2. Left hemisphere 2-column laminar 50 µm pitch PEDOT:PSS electrode locations**. Orange electrode models indicated on the brain across participants. Blue electrodes are clinical leads.

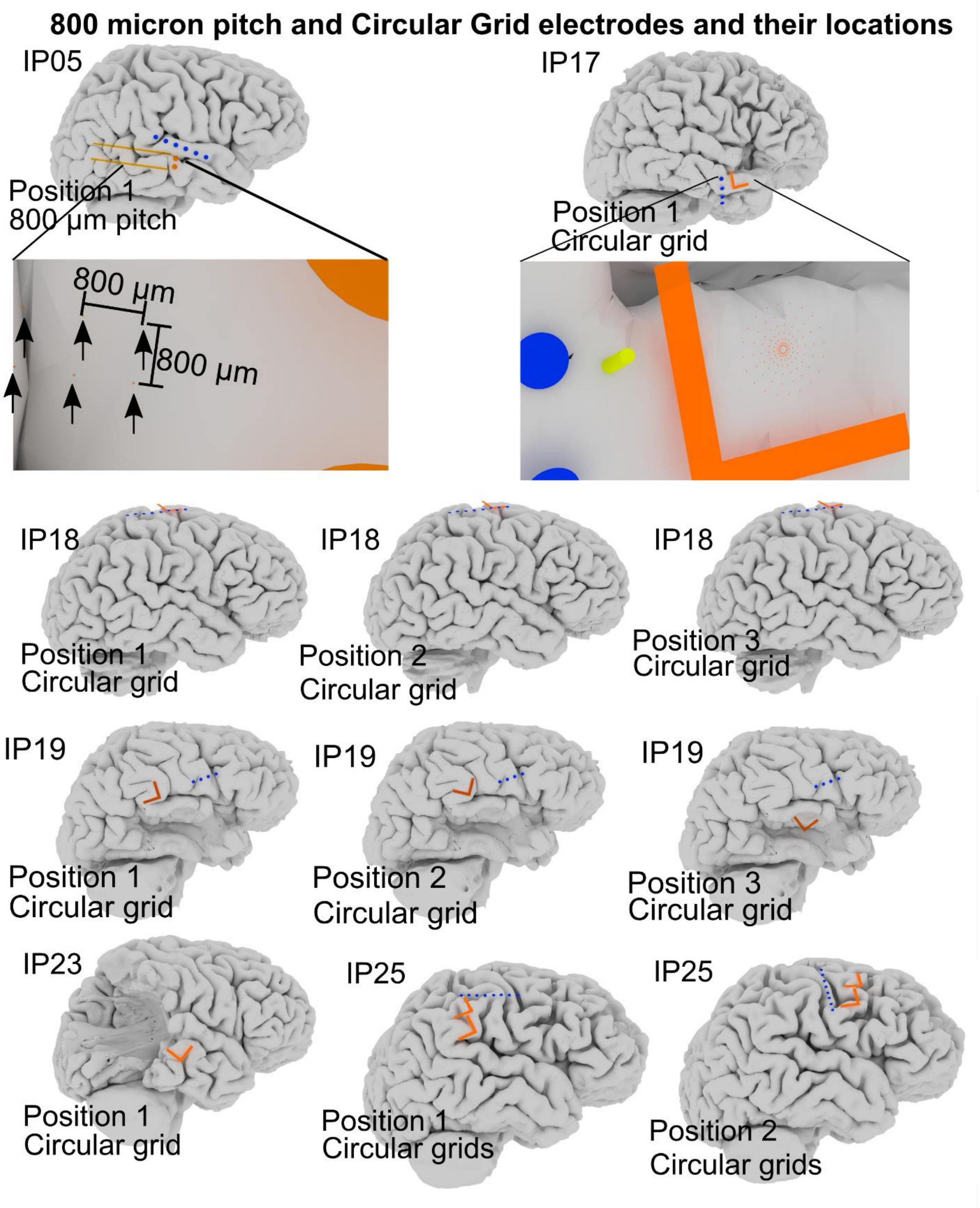

**Supplementary Fig. 3. PEDOT:PSS electrode locations of the 800 µm pitch 2-column laminar and the circular grid electrodes.** Orange electrode models indicated on the brain across participants. Blue electrodes are clinical leads.

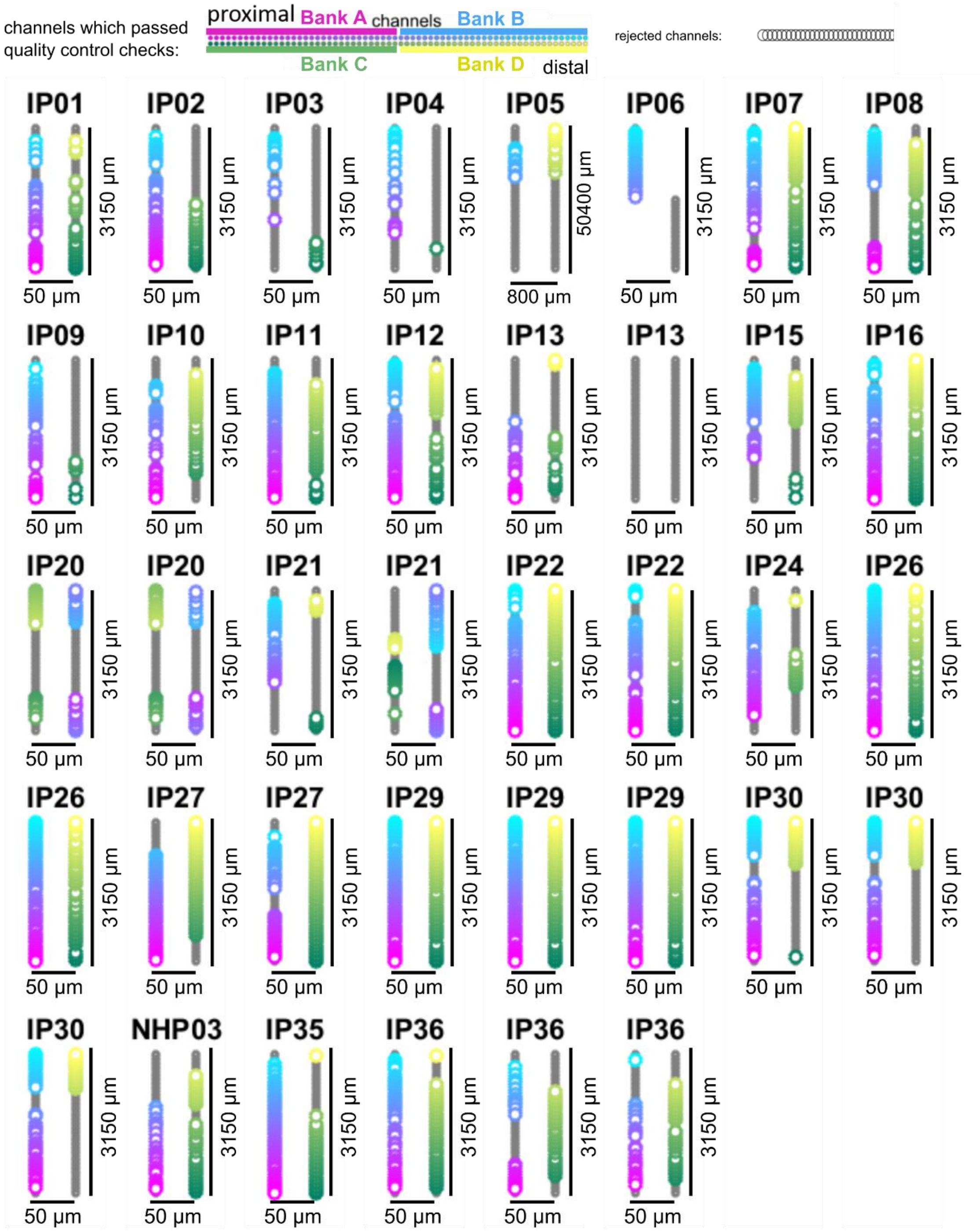

**Supplementary Fig. 4. Locations of electrode channels per electrode which survived quality control checks, 2-column grid.** The color coding of the waveforms match the color coding of the channels with magenta, yellow, green, and cyan color coding indicating the four separate 32 channel amplifier banks used in the recording. The color coding is cyan to magenta (channels 1-64) then from green to yellow (channels 65-128).

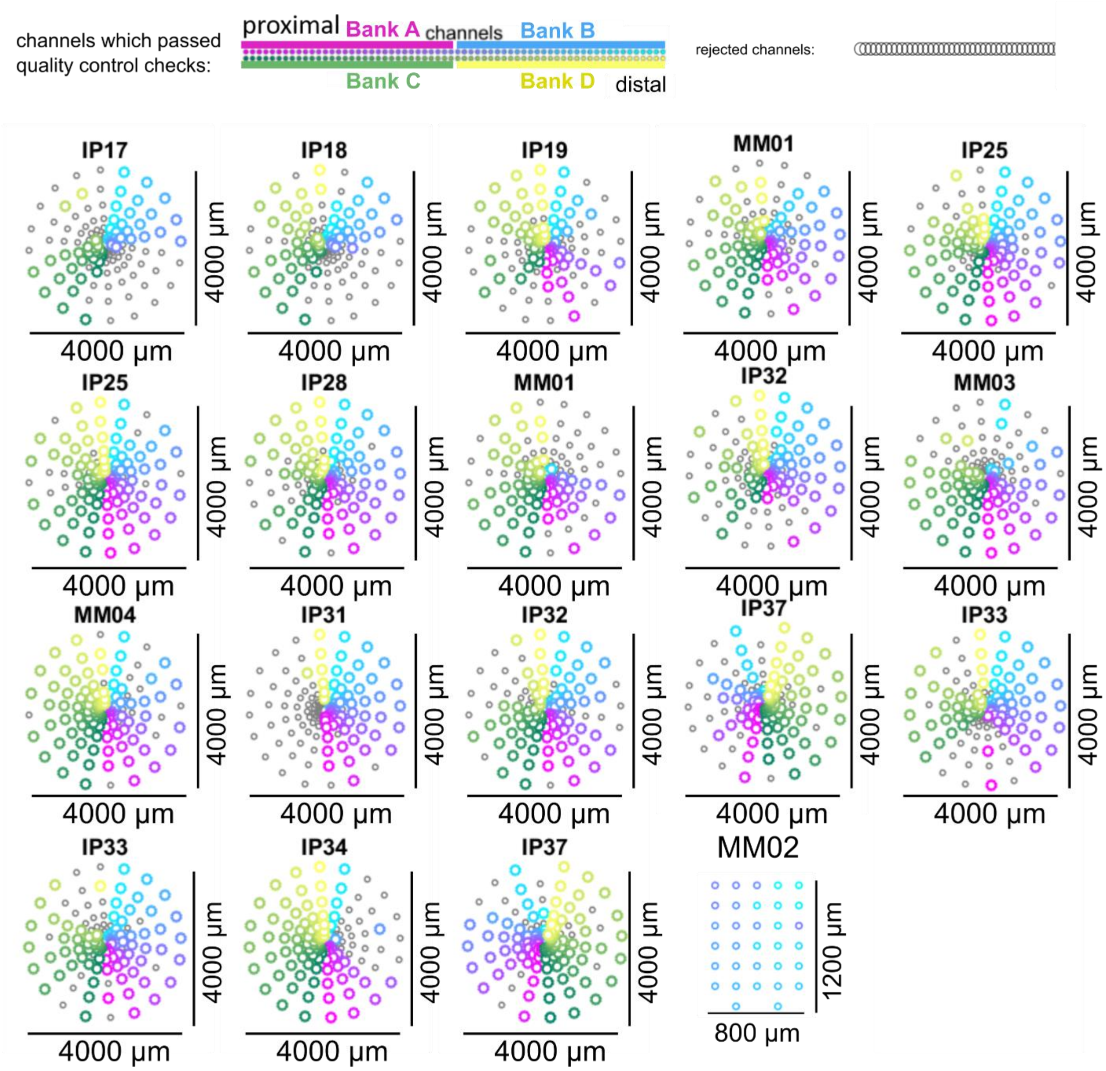

**Supplementary Fig. 5. Locations of electrode channels per electrode which survived quality control checks, Circular and rectangular grids**. The color coding of the waveforms match the color coding of the channels with magenta, yellow, green, and cyan color coding indicating the four separate 32 channel amplifier banks used in the recording. The color coding is cyan to magenta (channels 1-64) then from green to yellow (channels 65-128).

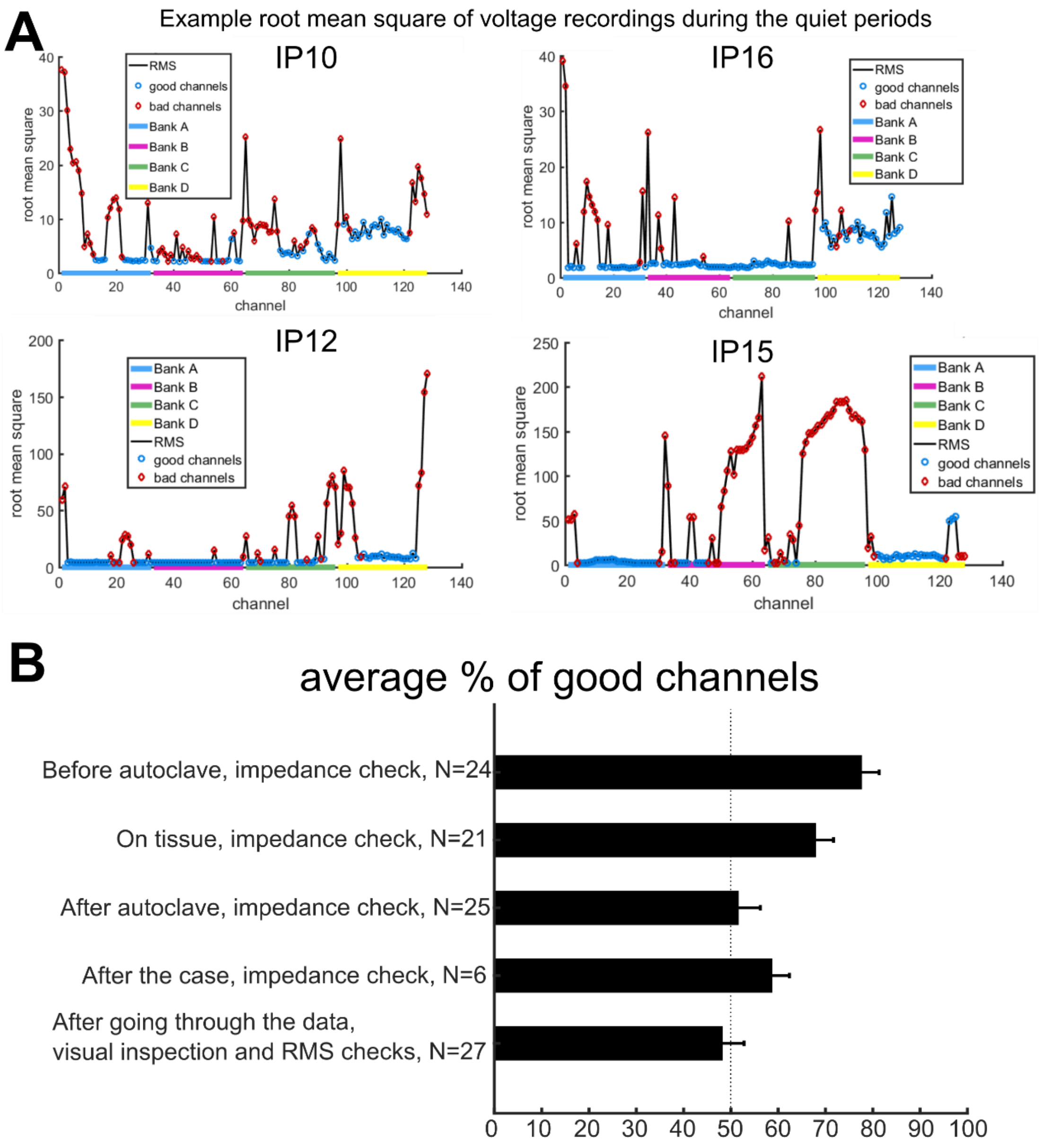

**Supplementary Fig. 6. Example RMS examination of channel activity during the quiet periods of a recording and impedance changes of the PEDOT:PSSelectrodes during each conditional step. A.** Example root mean square (RMS) of voltage recordings during quiet periods. Color coding indicate good or bad channels or which recording bank used in the testing. **B.** Average percentage of good channels after a series of different steps including sterilization and when on tissue.

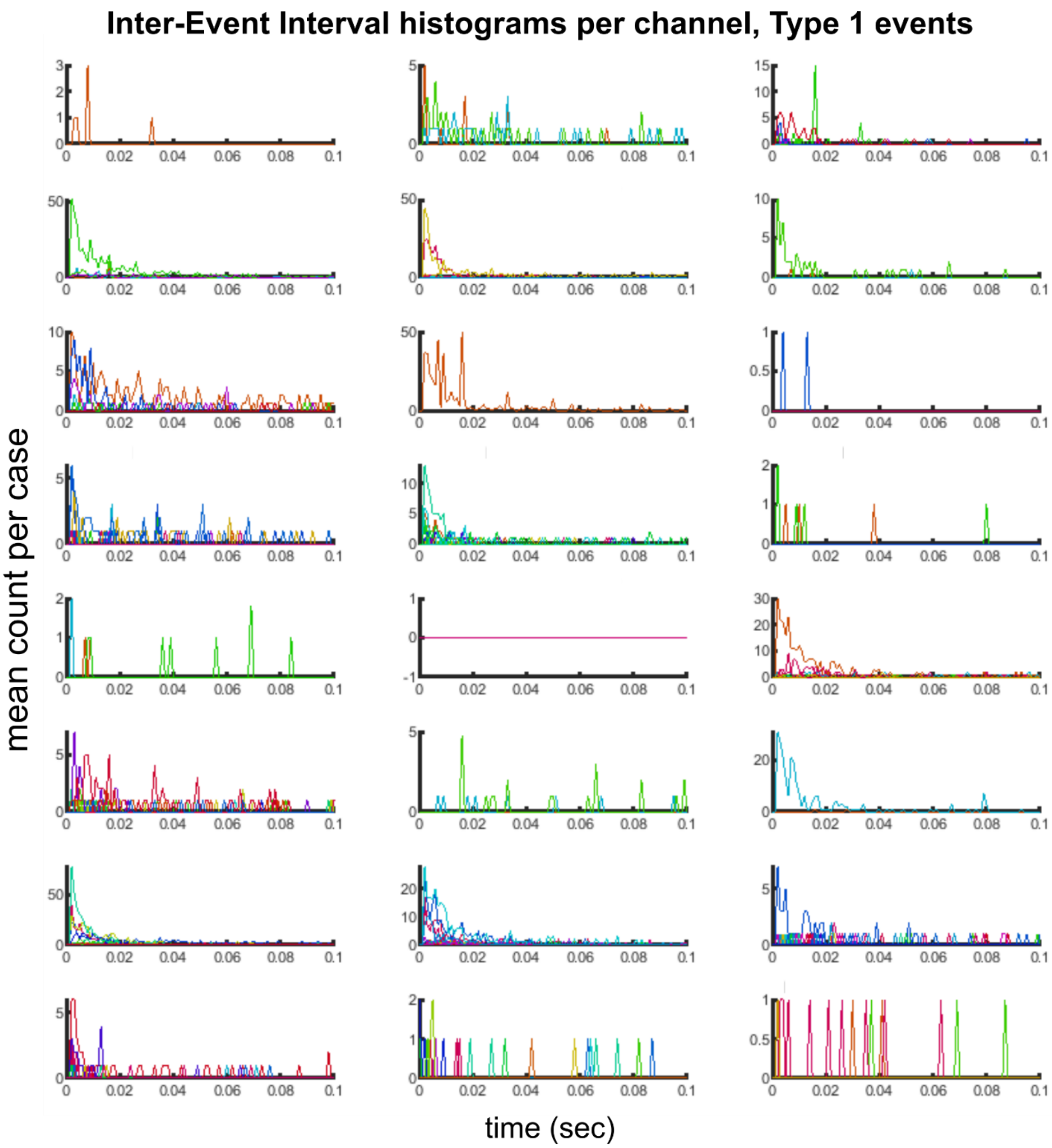

**Supplementary Fig. 7. Inter-Event-Interval (IEI) distribution for the Type 1 events.** Distribution of IEI values less than 0.1 sec. Each subplot is a different case, with individually differently colored lines indicate different clusters.

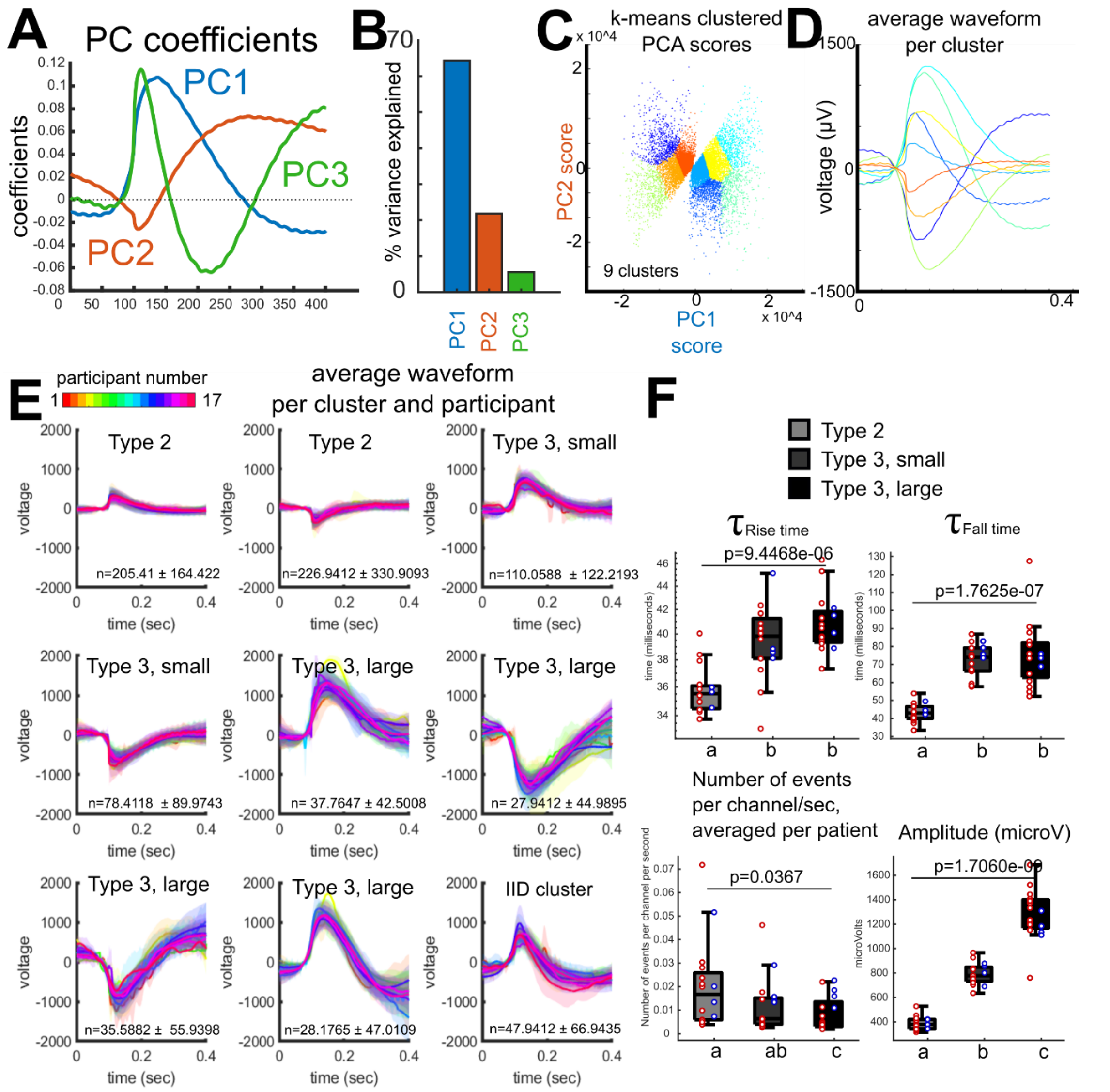

**Supplementary Fig. 8. Principal components analysis (PCA) on voltage waveforms to determine if there were different clusters of waveforms. A.** PCA coefficients applied to the entire data set, with the main waveforms identified as principal component 1 (PC1), principal component 2 (PC2), and principal component 3 (PC3). **B**. Percent variance explained by each principal component. **C**. k-means clustering of the PC1 vs PC2 scores. **D**. Average coefficient waveforms per cluster, color coded to the clusters shown in **C**. **E**. Average waveforms per participant in a subset of participants (N=17), showing different positive or negative deflections which are the basis for the template matched Type 2 or Type 3 events as well as IIDs. n-values indicate the number of events ± stdev. detected per participant per cluster. Averages are shown with shaded areas reflecting s.e.m. **F**. Average $\tau_{\text{rise time}}$, $\tau_{\text{fall time}}$, frequency of events, and absolute amplitude per waveform type across participants, shown in box plots with confidence bounds. N=17. p-values are for comparisons between waveform types, Wilcoxon rank-sum comparison. The waveforms and numbers here are not the exact same as the template-matched waveforms as in the remainder of the study as this sorting was to determine the distribution and types of diverse waveforms possible using a threshold-crossing detection.

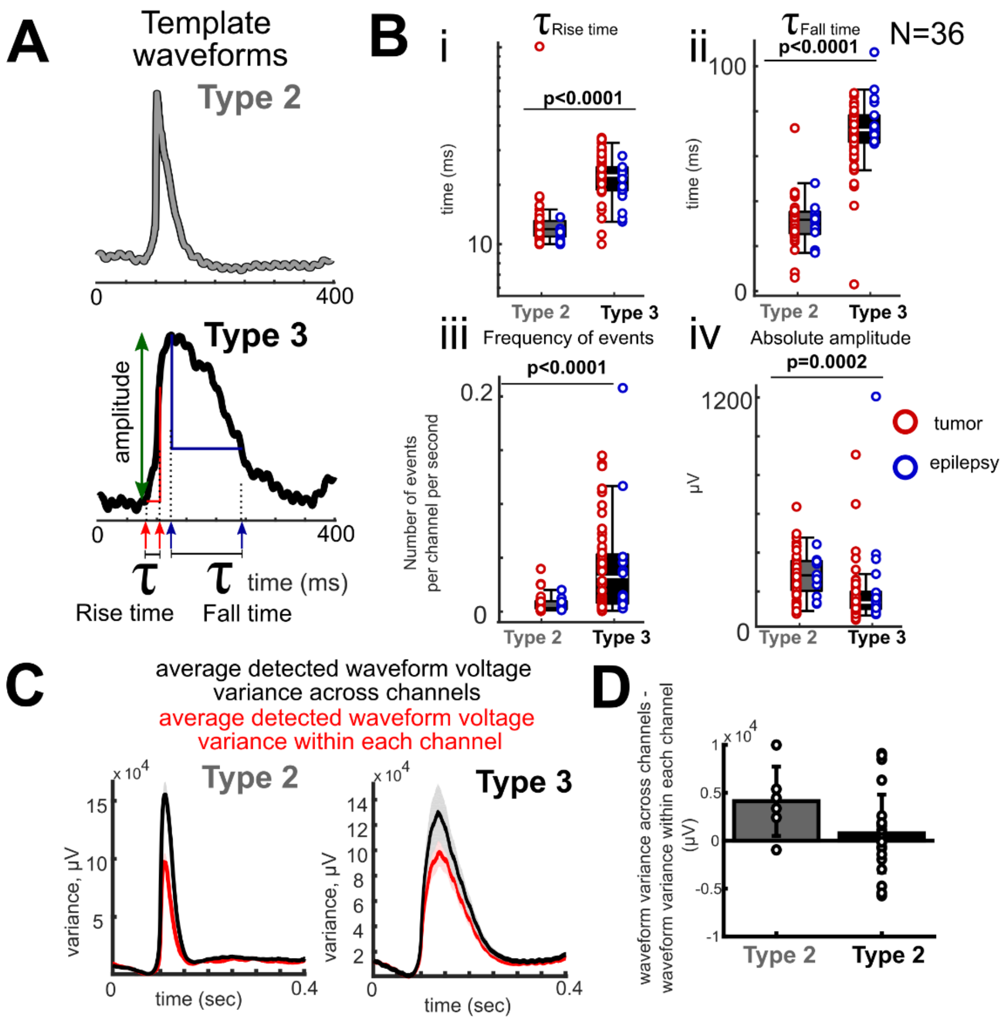

**Supplementary Figure 9. Criteria for detecting Type 2 and 3 events and variation across channels versus within channels**. **A**. Template waveforms used to classify the activity. Measurements include baseline activity before the onset to the peak amplitude (green arrow), the rise and fall times relative to the peaks (see methods) by quantifying the $\tau_{\text{rise time}}$ and $\tau_{\text{fall time}}$ time constants as in the following equation: $\Delta V = (V_{final} - V_{start})*(1-1/e^{\wedge}(time/\ \tau))$, where $\tau_{\text{rise time}}$ was calculated as time it took for the voltage to reach 0.63 of the maximum peak (red arrows) and $\tau_{\text{fall time}}$ was calculated as the time it took for the voltage to reach 0.37 of the fall back to baseline (blue arrows). **B**. Average $\tau_{\text{rise time}}$ (**Bi**), $\tau_{\text{fall time}}$ (**Bii**), frequency of events (**Biii**), and absolute amplitude (**Biv**) per waveform type across participants, shown in box plots with confidence bounds. N=36. p-values are for comparisons between waveform types, Wilcoxon rank-sum comparison. Center lines indicate median, box limits indicate confidence bounds. **C**. Left: Average variance of the voltage for detected Type 2 events per channel (red line) and the average variance of the voltage for detected Type 2 events across channels (black line) per recording and averaged across all participants (N=29). Shaded regions indicate s.e.m. Right: Average variance of the voltage for detected Type 3 events per channel (red line) and the average variance of the voltage for detected Type 3 events across channels (black line) per recording and averaged across all participants (N=29). Shaded regions indicate s.e.m. **E**. Difference between waveform variance across channels and the waveform variance within each channel, averaged across all participants (N=36). Comparisons are only made for instances when more than 10 waveforms were detected during baseline activity per channel.

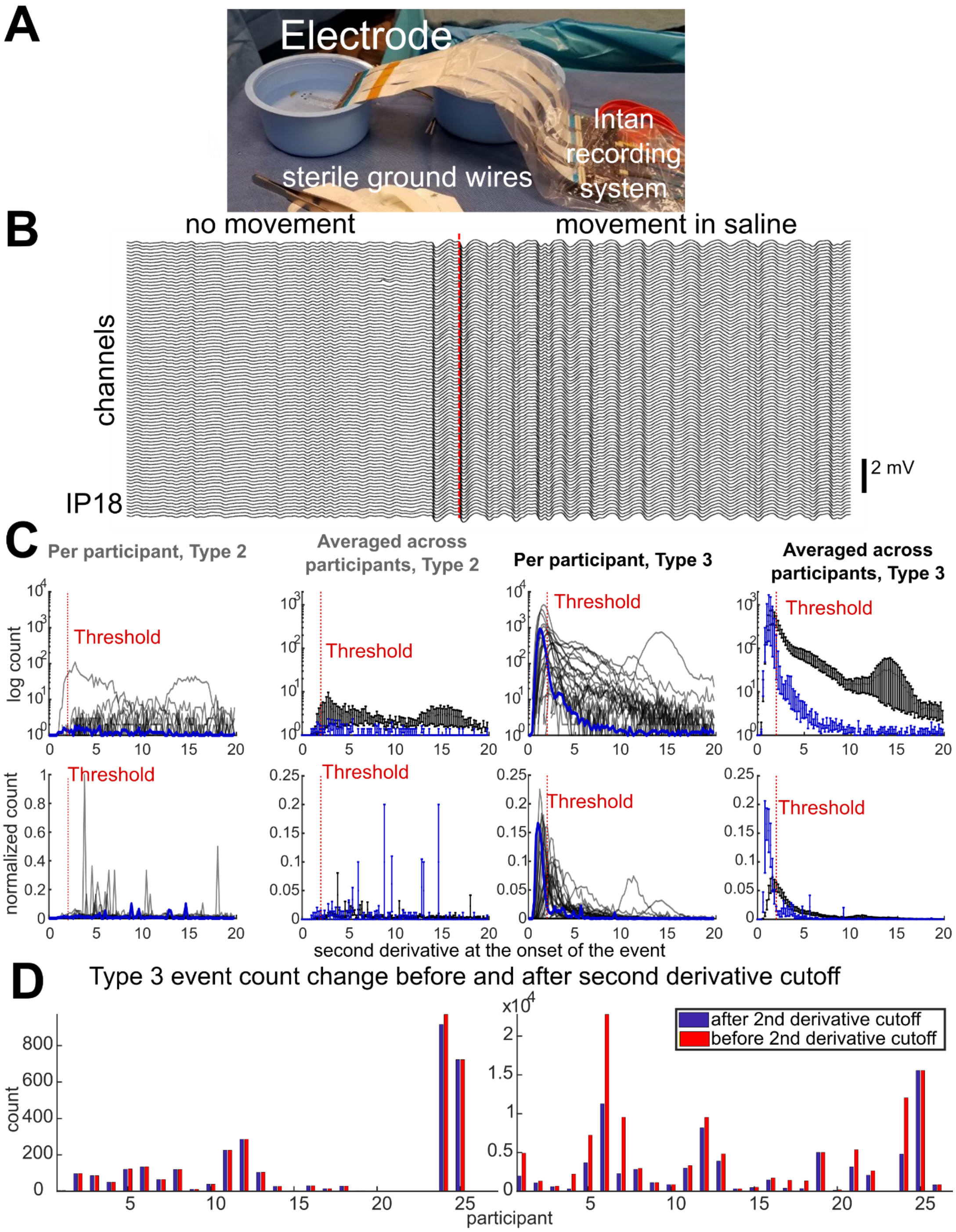

**Supplementary Fig. 10. Detecting and separating Type 2 and Type 3 waveforms in saline recordings versus on-brain recordings using the same electrodes in the operating room**. **A**. Example recording set up. **B**. Example recording with induced movement of the electrode in saline. **C**. Top: log count of the second derivative of the first 100 ms of the waveform in the saline (blue) and over brain (black lines) for the Type 2 (two left plots) and Type 3 (two right plots) for individual and averaged values.

Bottom: normalized count (normalized by the total number of events) the second derivative of the first 100 ms of the waveform in the saline (blue) and over brain (black lines) for the Type 2 (two left plots) and Type 3 (two right plots) for individual and averaged values. The red lines indicate the cutoff threshold used to remove waveforms from the on-brain recordings for further analyses. **D**. Number of events removed per participant and recording before and after added second derivative thresholding approach.

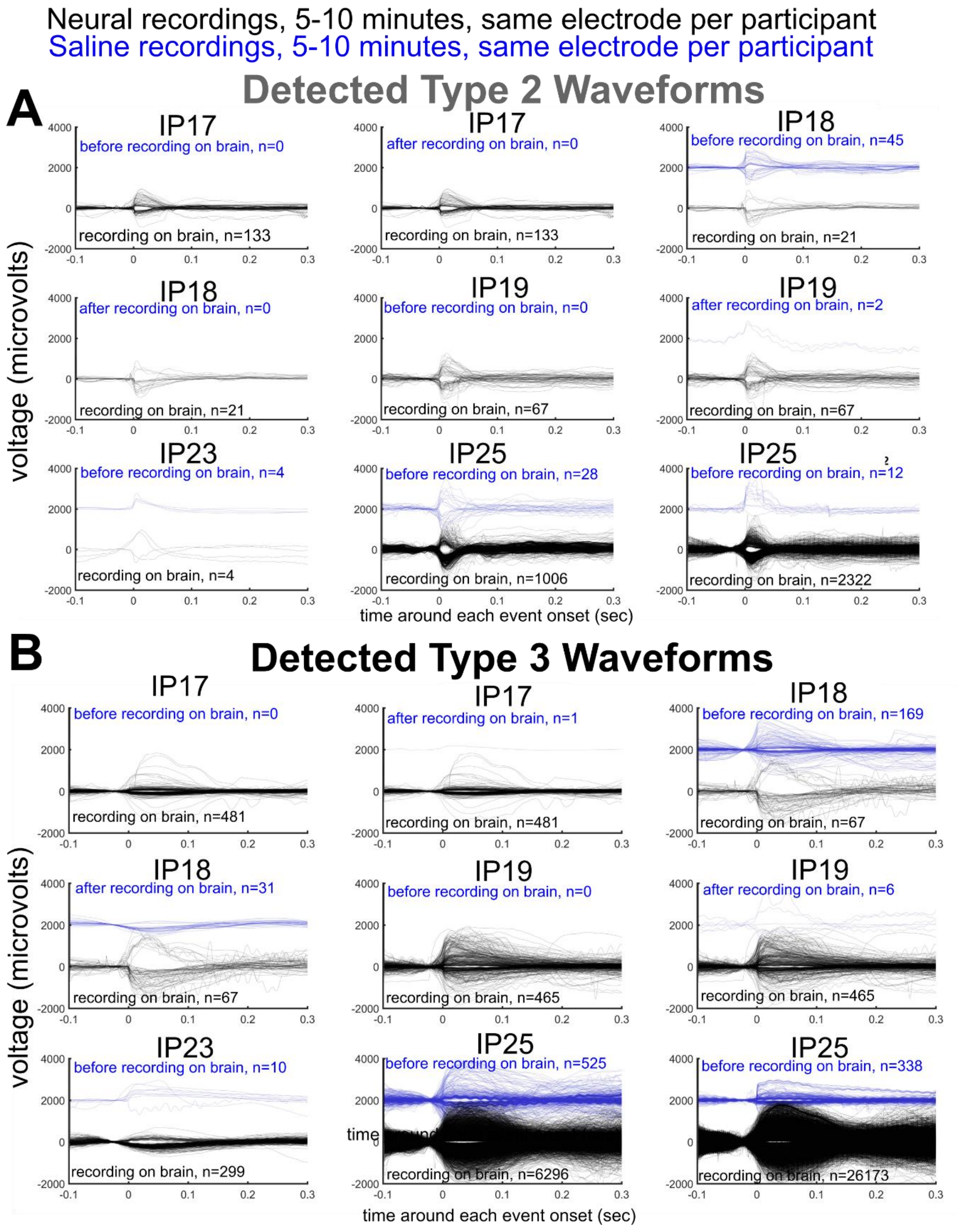

**Supplementary Fig. 11. Type 2 and Type 3 waveforms in saline recordings versus on-brain recordings using the same electrodes in the operating room**. **A**. Detected Type 2 waveforms with the same electrodes over brain (black lines) or in saline (blue lines). **B**. Detected Type 3 waveforms with the same electrodes over brain (black lines) or in saline (blue lines).

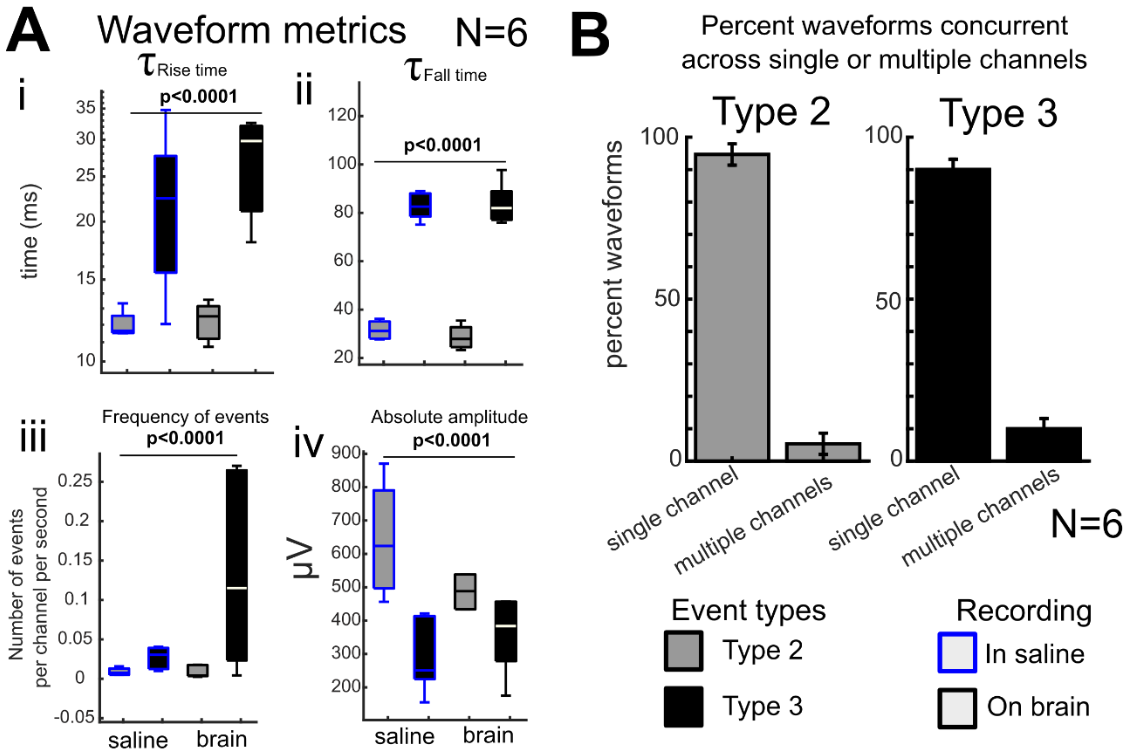

**Supplementary Fig. 12. Saline recording waveform metrics compared to recordings on brain with the same sets of electrodes**. Average $\tau_{\text{rise time}}$ (**Ai**), $\tau_{\text{fall time}}$ (**Aii**), inter-event interval (**Aiii**), and Absolute amplitude (**Aiv**) per waveform type across Type 2 and Type 3 events in saline (blue outlined boxplots) and over brain (black outlined boxplots) with the same electrodes (N=6). p-values are the results of Wilcoxon rank-sum tests comparing Type 2 and 3 events per type and waveform measurement and condition. N=6 participants or instances of testing. **B**. Percent of waveforms detected across single or multiple channels during saline recordings.

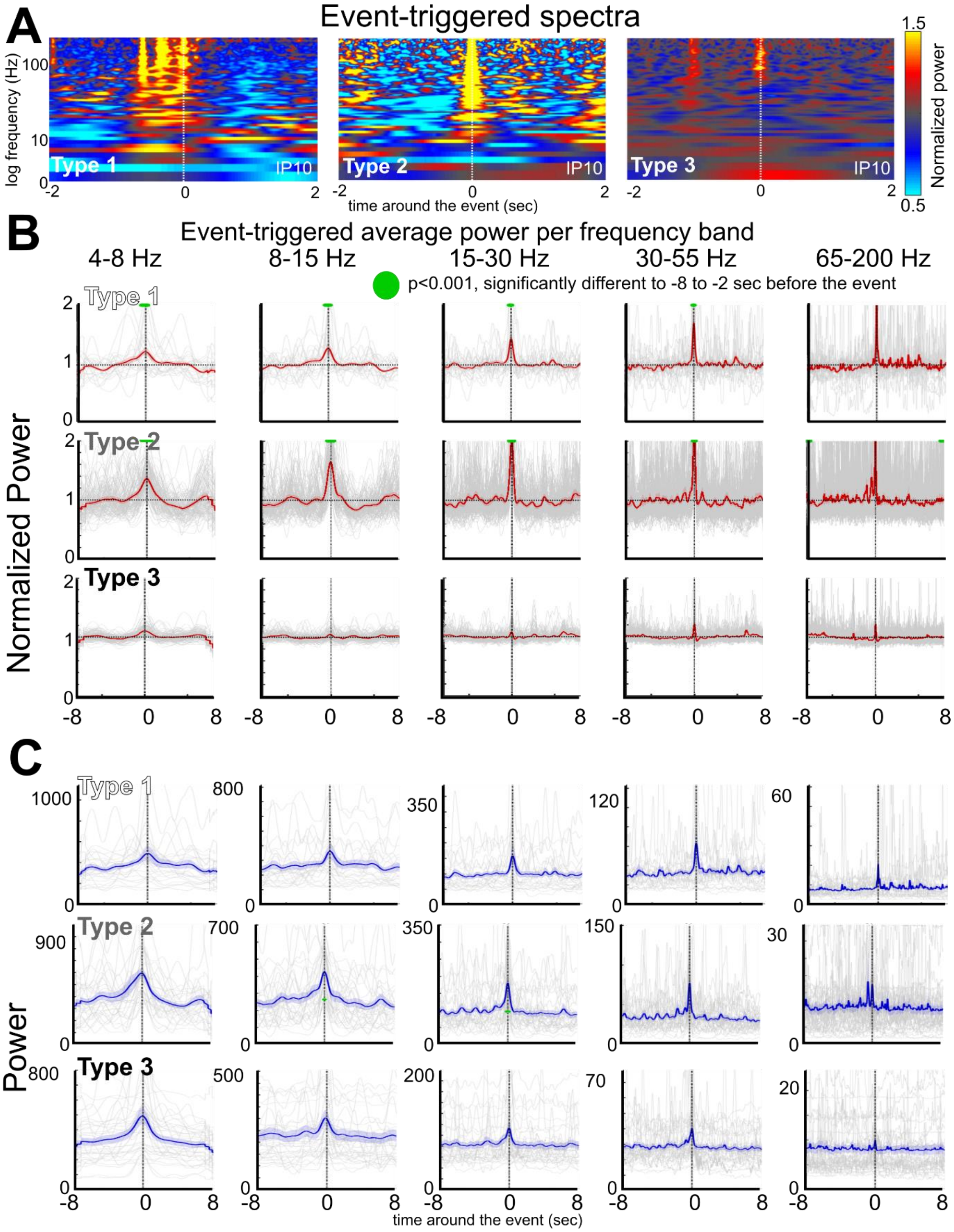

**Supplementary Fig. 13. Event-triggered local field potential (LFP) and spectral changes**. **A**. Average power spectra aligned to Type 1 (left), Type 2 (middle), and Type 3 (right) events normalized to the entire recording per frequency band (N=1). **B**. Average normalized power along a sliding moving window (moved every 10 ms, normalized to the entire recording per frequency band) shown as a red line aligned to the onset of the detected Type 1 (top), Type 2 (middle) and Type 3 (bottom) events, ± 8 seconds. Grey lines are individual recordings, N=29 participants. Shaded area indicates s.e.m. Green dots indicate time steps when the power was significantly different to baseline, which is between 1 and 3 seconds before the event, $p<0.001$, Wilcoxon rank sum test, false discovery rate controlled for multiple comparisons. **C**. Average power along a sliding moving window (moved every 10 ms, normalized to the entire recording per frequency band) shown as a blue line aligned to the onset of the detected Type 1 (top), Type 2 (middle) and Type 3 (bottom) events, ± 8 seconds. Shaded area indicates s.e.m. Grey lines are individual recordings, N=29 participants.

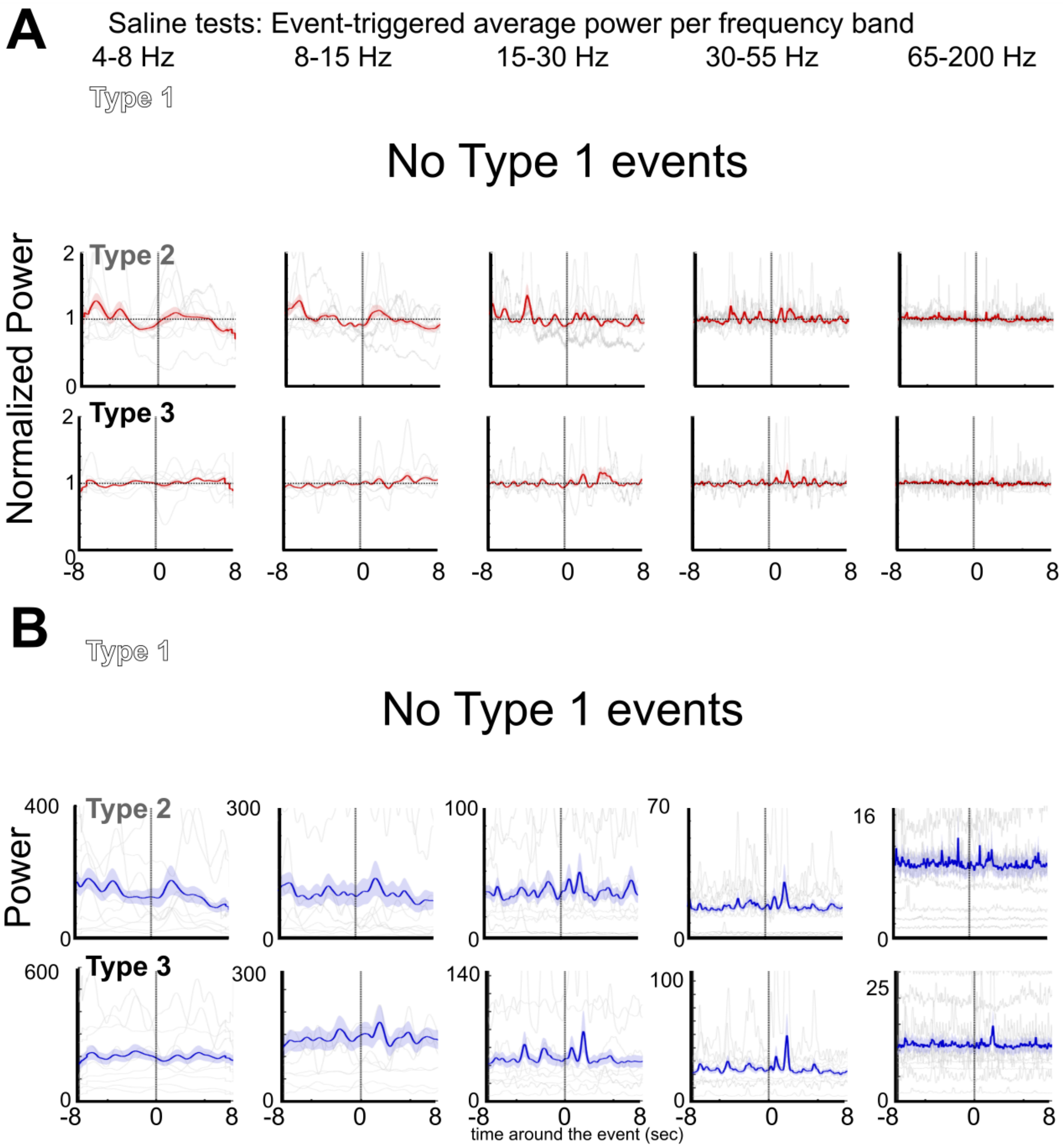

**Supplementary Fig. 14. Event-triggered time-locked power dynamics, saline**. **A**. Average normalized power along a sliding moving window (moved every 10 ms, normalized to the entire recording per frequency band) shown as a red line aligned to the onset of the detected Type 2 (middle) and Type 3 (bottom) events, ± 8 seconds. Grey lines are individual recordings, N=4 mice, N=1 NHP. Shaded area indicates s.e.m. **B**. Average power along a sliding moving window (moved every 10 ms, normalized to the entire recording per frequency band) shown as a blue line aligned to the onset of the detected Type 2 (middle) and Type 3 (bottom) events, ± 8 seconds. Shaded area indicates s.e.m. Grey lines are individual recordings.

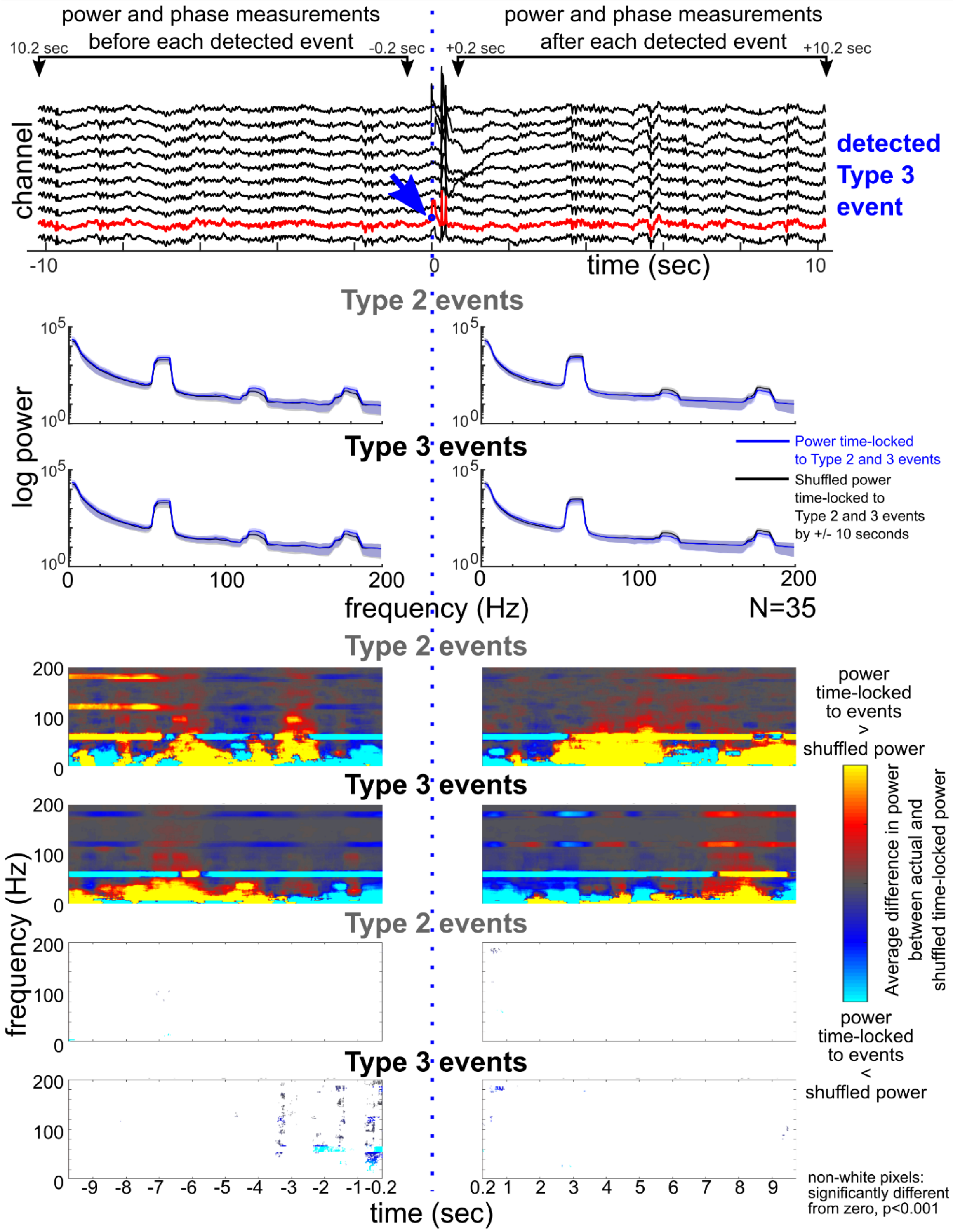

**Supplementary Fig. 15. Power in different frequency bands surrounding Type 2 and 3 events**. **A**. Example traces with an event detected aligned to zero. The windows of time for analysis ± 10 sec around the event. Then, the steps involved the following: a) calculating spectrograms 10 sec before and after the Type 2/3 events (not including+/-

200 ms around each event). b) performing the same calculation but, instead of centering it around the Type 2/3 events, add or subtract a time in the range of +/- 10 seconds (jittered time-locked power calculation). **B**. The power curves across frequencies before and after the events and in the time-jittered power calculations for the Type 2 and Type 3 events. **C**. Difference in the spectrograms across frequencies before and after the events for the time-locked spectrograms versus the time-jittered power calculations for the Type 2 and Type 3 events on the per-channel basis, averaged across participants. **D**. Plotting significant values if the time-jittered versus time-locked value differences are significantly different from zero across the data set. Wilcoxon rank-sum test for significance.

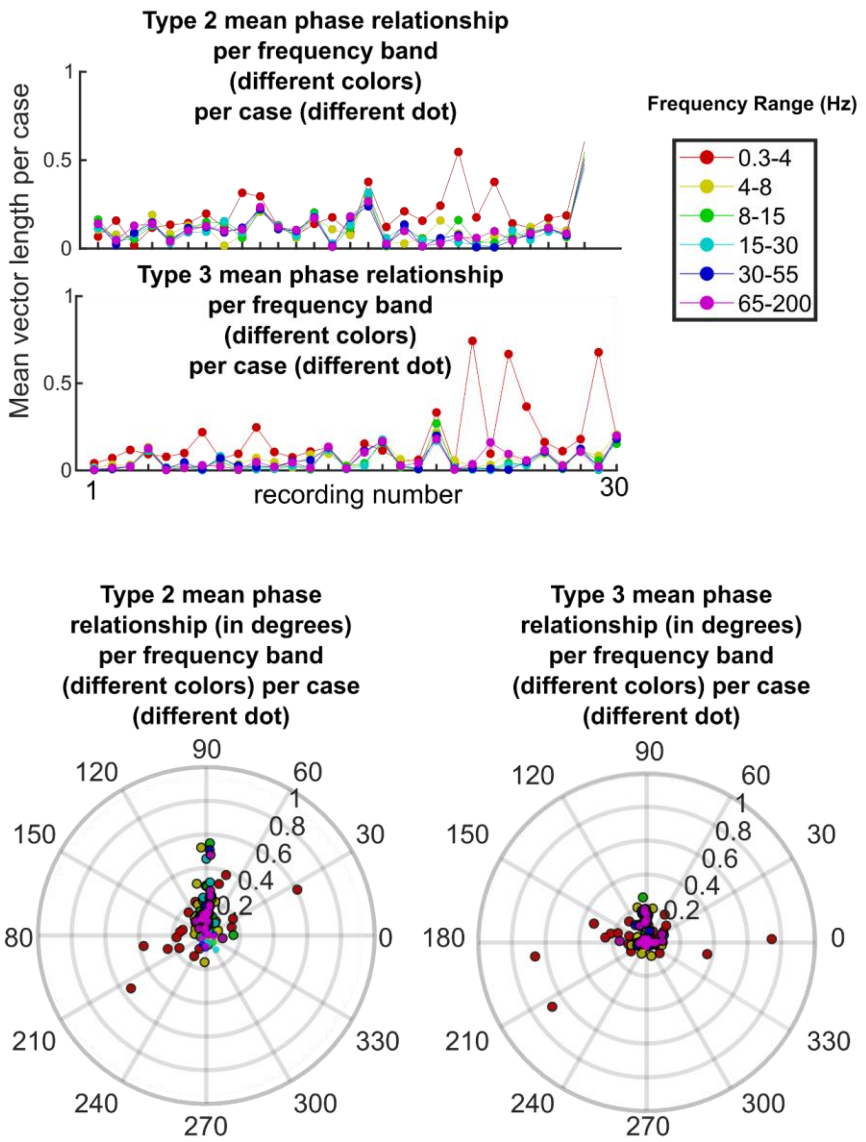

**Supplementary Fig. 16. Testing phase-locking per Type 2 and 3 event per frequency band**. **A**. To calculate the phase relationships per frequency band, we filtered the signal to each frequency band with eegfilt (EEGLab package; (37)) and then calculated phase per frequency band (Hilbert transform) and averaged those phase values using the CircStat toolbox (Berens, 2009). As we were performing these calculations on a per-channel and then per-case basis, we had to average phase values when the number of events per recording was greater than 5 as phase is highly susceptible to low event counts. This resulted in calculated average phase angles and lengths of the vectors per case after averaging these values across channels for each of 6 frequency bands (**A**). **B**. We also calculated the average phase angles for each of 6 frequency bands for the Type 2 and 3 events.

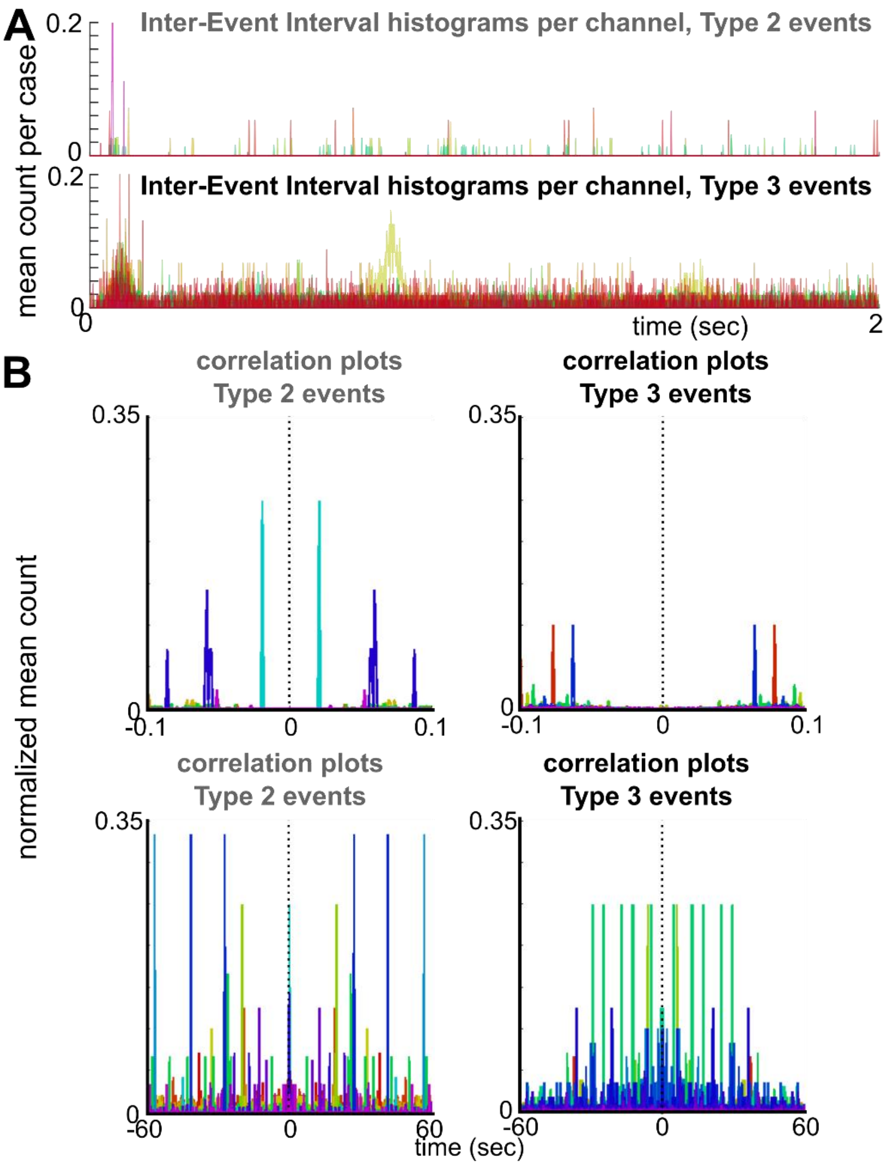

**Supplementary Fig. 17. Inter-Event-Interval (IEI) and autocorrelation dynamics for the Type 2 and Type 3 events. A.** Average per-case Inter-event Interval histograms per channel for the Type 2 (top) and Type 3 (bottom) events. Each line is a different case. The majority of the cases do not have IEI values less than 2 seconds while a few cases demonstrate IEI events below than 2 sec. **B.** Cross-correlation performed by cross-correlating single events with all the other events per channel per Type 2 (left) and Type 3 (right) events at two different time scales from -0.1 to 0.1 sec (top) and from -60 to 60 sec. Each color line is a different case. While some cases have some type of regularity in the cross-correlation, there was not any obvious rhythmic consistency across cases or even within a case.

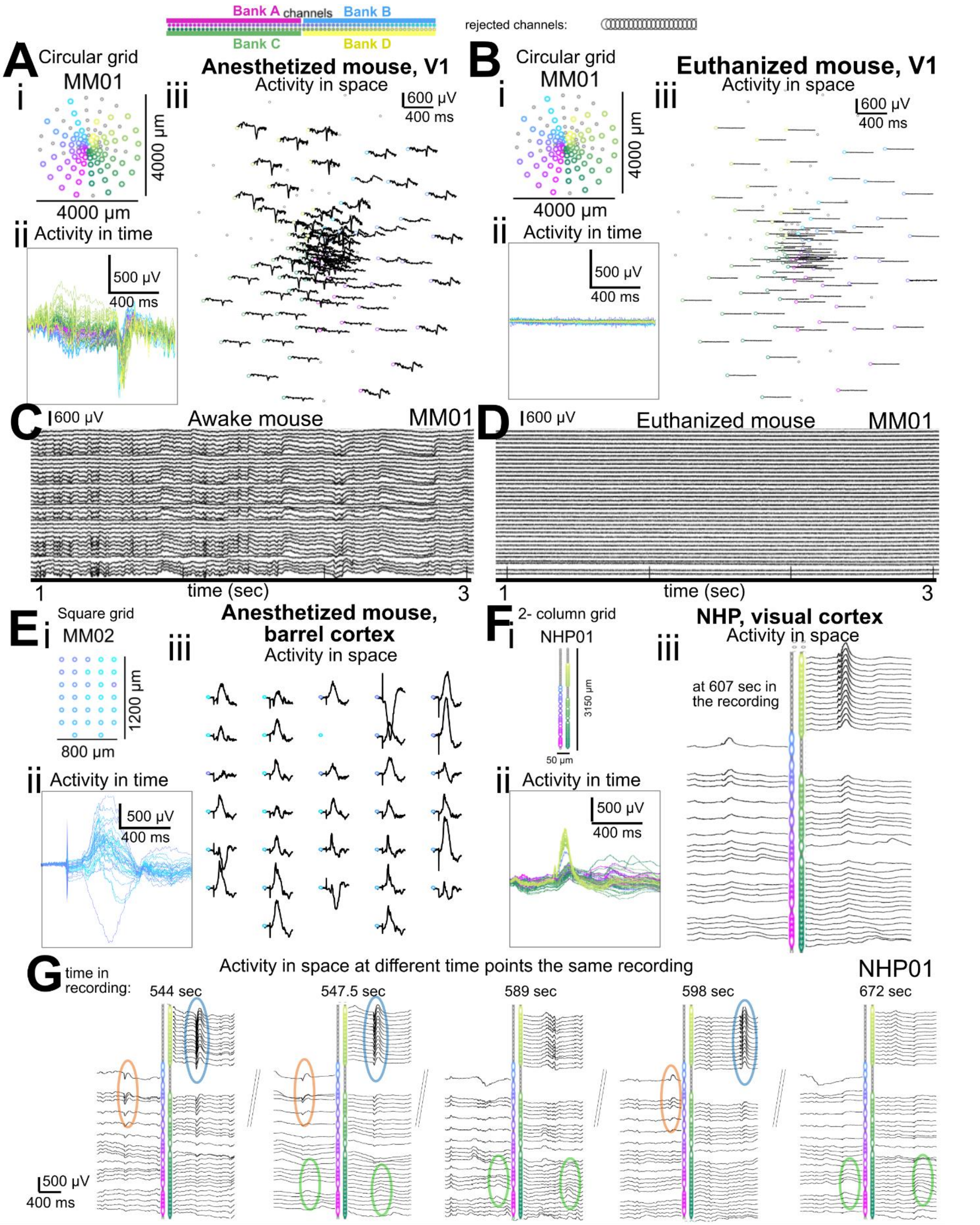

**Supplementary Fig. 18. NHP and mouse recordings. A**. Example recording (electrodes which passed criteria, **Ai**) from mouse V1 (in a ketamine-anesthetized mouse) showing activity changes in time (**Aii**) and a proportion of electrodes and their activity mapped to the two-column electrode array (**Aiii**). **B**. Recording from the same mouse after euthanazia in time (**Bi-ii**) and mapped to the circular grid (**Biii**). The color coding of the waveforms match the color coding of the channels with magenta, yellow, green, and cyan color coding indicating the four separate 32 channel amplifier banks used in the recording. **E**. Example recording from a square grid (electrodes which passed criteria, **Ci**) from mouse barrel cortex (in an anesthetized mouse) showing activity changes in time (**Cii**) and a proportion of electrodes and their activity mapped to the two-column electrode array (**Ciii**). **F**. Example recording from a 2-column grid (electrodes which passed criteria, **Fi**) from NHP visual cortex (in an anesthetized NHP) showing activity changes in time (**Fii**) and a proportion of electrodes and their activity mapped to the two-column electrode array (**Fiii**). **G**. Example recordings of events at different time points (for the NHP) showing only a subset of electrode channels, with repeated events shown through time.

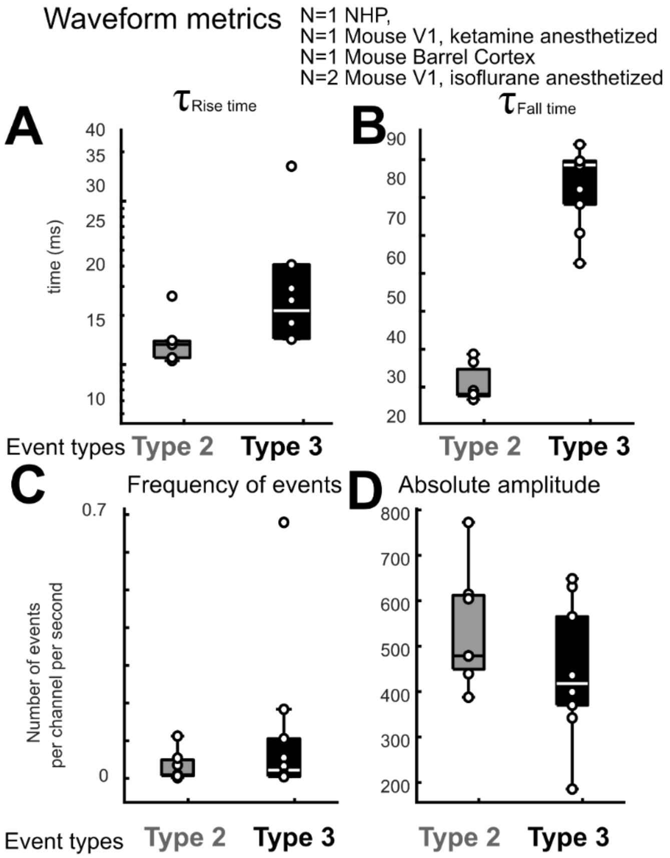

**Supplementary Fig. 19. NHP and mouse recordings, waveform characteristics.** Average $\tau_{\text{rise time}}$ (**A**), $\tau_{\text{fall time}}$ (**B**), inter-event interval (**C**), and Absolute amplitude (**D**) per waveform type across Type 2 and Type 3 events across NHP and mouse recordings. p-values are the results of Wilcoxon rank-sum tests comparing Type 2 and 3 events per type and waveform measurement. N=4 mice, N=1 NHP.

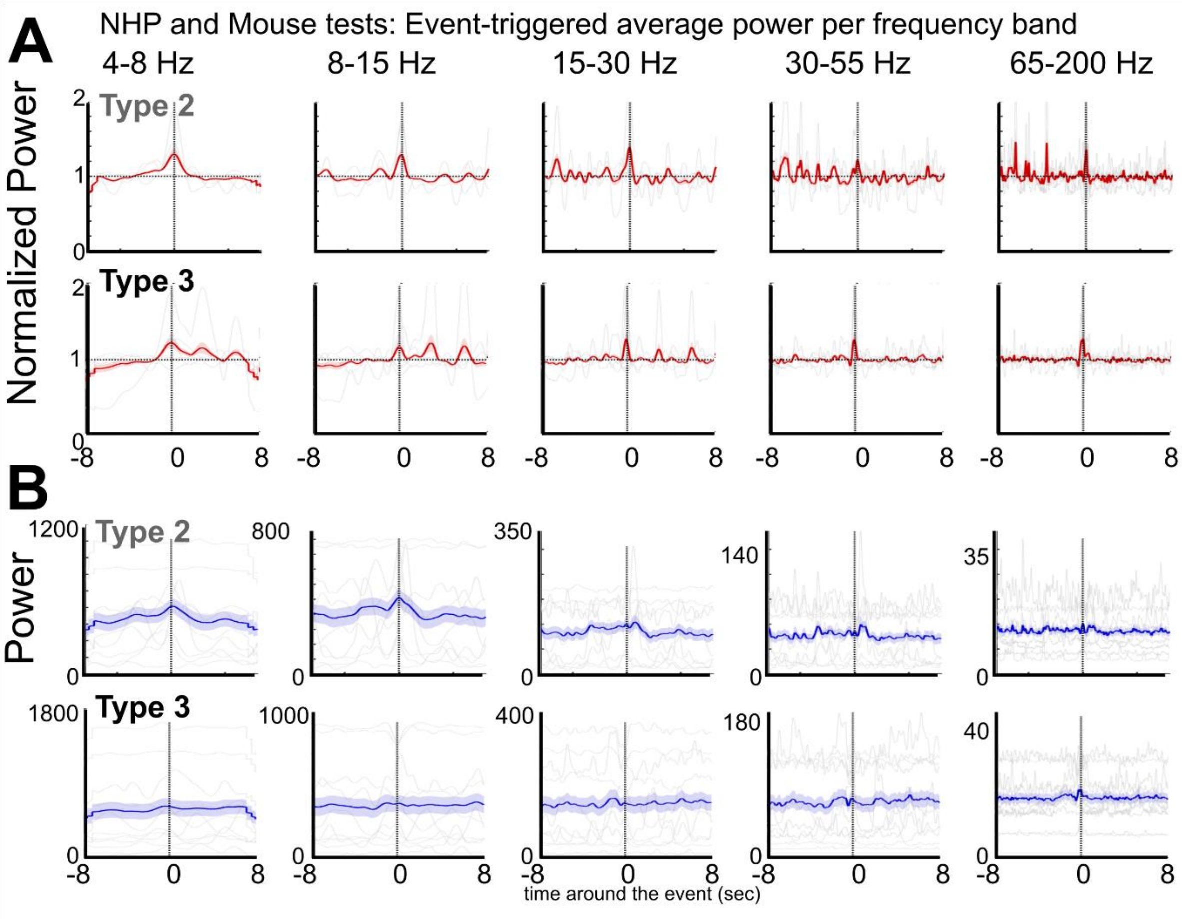

**Supplementary Fig. 20. Event-triggered time-locked power dynamics for Type 2 and 3 events across NHP and mouse recordings**. **A**. Average normalized power along a sliding moving window (moved every 10 ms, normalized to the entire recording per frequency band) shown as a red line aligned to the onset of the detected Type 1 (top), Type 2 (middle) and Type 3 (bottom) events, ± 8 seconds. Grey lines are individual recordings, N=4 mice, N=1 NHP. Shaded area indicates s.e.m. **B**. Average power along a sliding moving window (moved every 10 ms, normalized to the entire recording per frequency band) shown as a blue line aligned to the onset of the detected Type 1 (top), Type 2 (middle) and Type 3 (bottom) events, ± 8 seconds. Shaded area indicates s.e.m. Grey lines are individual recordings, N=4 mice, N=1 NHP.

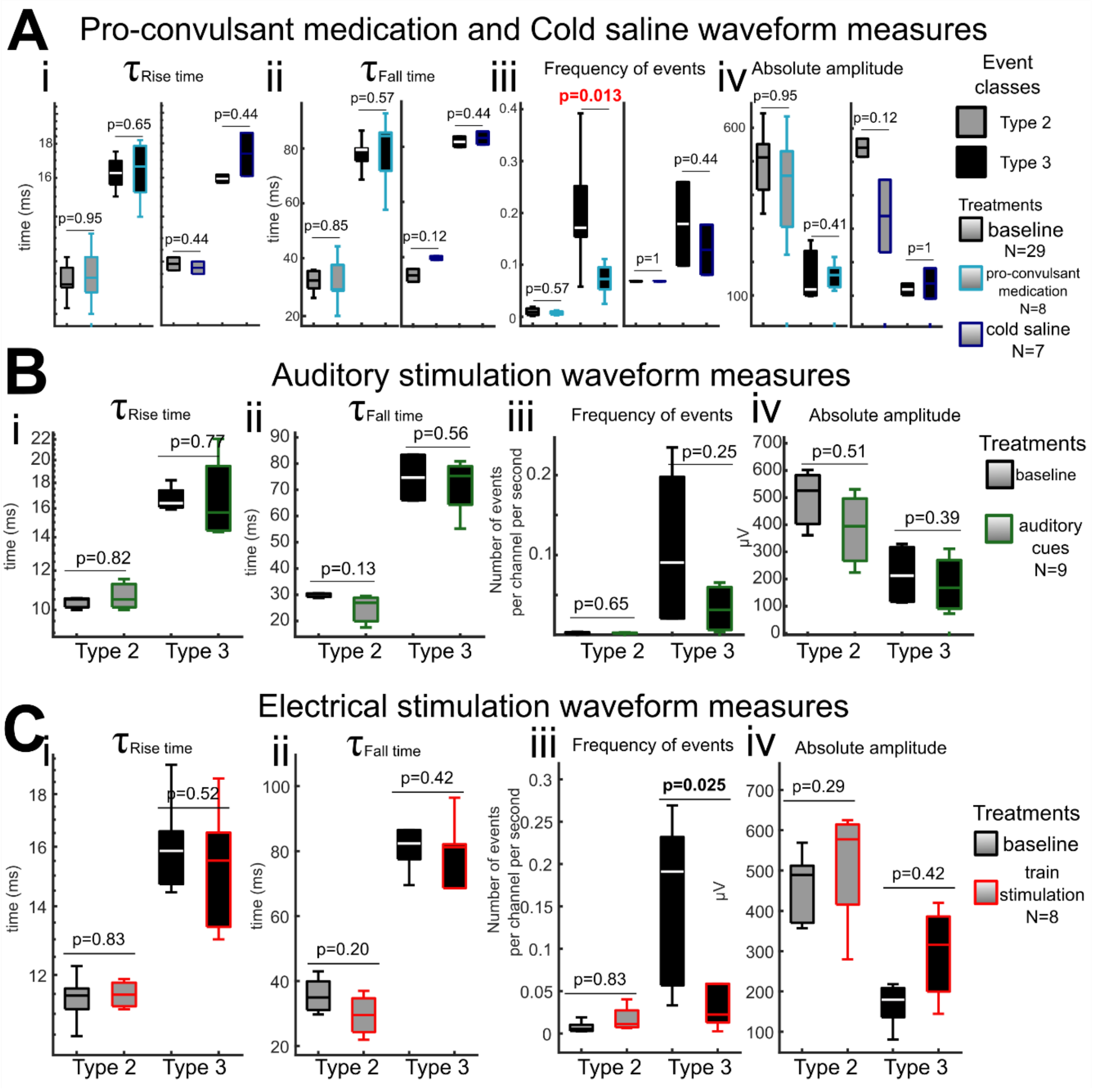

**Supplementary Fig. 21. Waveform measurements during the application of pro-convulsant medication, cold saline, auditory stimulation and electrical stimulation. A-C**. Average $\tau_{\text{rise time}}$ (**i**), $\tau_{\text{fall time}}$ (**ii**), inter-event interval (**iii**), and Absolute amplitude (**iv**) per waveform type across participants for the different physiological treatments. p-values are the results of Wilcoxon rank-sum tests comparing baseline events to the minutes after the manipulation (orange brackets in **Fig. 4C** (for **A**), **5C** (for **B**), and **6C** (for **C**)) per type and waveform measurement. N-values indicated in **A-C**.

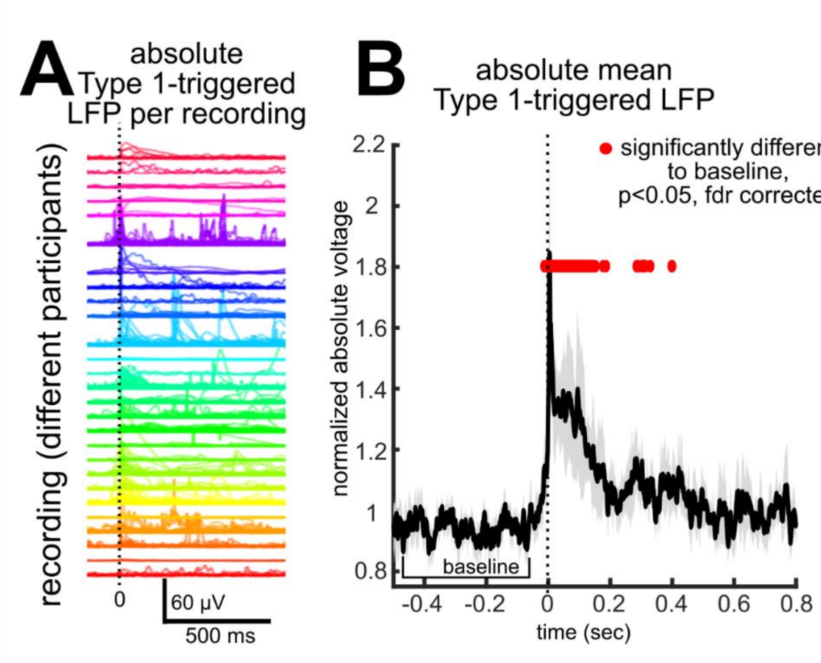

**Supplementary Fig. 22. Spike-triggered voltage averages around Type 1 events. A.** Spike-triggered voltage averages of the low frequency (LFP) data per recording across multiple events**.** Each line and color indicate a different recording across N=29 participants. **B**. Averaged normalized (to baseline) LFP voltage deflection around Type 1 events, N=29.
